# Diversity and evolution of the ribovirian class *Stelpaviricetes*

**DOI:** 10.64898/2026.08.03.742603

**Authors:** Anastasia Gulyaeva, Pascal Mutz, Qiyan Liu, Yuri I. Wolf, Guan-Zhu Han, Eugene V. Koonin, Valerian V. Dolja

## Abstract

Metatranscriptome mining has dramatically expanded the known diversity of ribovirians at all taxonomic levels. We explored in detail the class *Stelpaviricetes* in the phylum *Pisuviricota* that so far included 3 orders and 6 recognized virus families, order *Stellavirales* with family *Astroviridae* infecting vertebrates, *Patatavirales* with families *Potyviridae,* the largest known family of plant ribovirians, and *Potyliviridae,* and *Hypofuvirales* consisting of 3 families of viruses associated with phytopathogenic fungi. Using an order-specific collection of Hidden Markov Model profiles for the RNA-dependent RNA polymerase (RdRP), the hallmark protein of ribovirians, we identified 103 putative families within *Stelpaviricetes,* most of which share a uniform, astrovirus-like genome architecture, encoding three recognizable protein domains, the RdRP, a chymotrypsin-like protease and a single jelly roll capsid protein. The topology of the phylogenetic tree of the RdRP implies that the common ancestor of *Stelpaviricetes* had an astrovirus-like genome. The ancestor of the order *Patatavirales* acquired two additional domains, a papain-like protease and a superfamily 2 helicase, whereas the capsid protein was replaced with an unrelated one forming filamentous capsids. The ancestor of *Hypofuvirales* also encoded a superfamily 2 helicase and, possibly, a papain-like protease, but lost the capsid protein. *Hypofuvirales* might have derived from *Patatavirales* or evolved independently from an astro-like ancestor. For the vast majority of *Stelpaviricetes* identified in metatranscriptomes, host assignment remains elusive. Nevertheless, analysis of endogenous virus elements combined with information on isolated viruses provides some clues, in particular, suggesting that the common ancestor of *Potyviridae* and *Potyliviridae* had a fungal host whereas the common ancestor of *Stelpaviricetes* might have been a protist virus.

**IMPORTANCE:** The enormous diversity of viruses on earth is only now coming to light through extensive metagenome and metatranscriptome mining. Using vast databases of ribovirus genomes, we performed phylogenomic analysis of the class *Stelpaviricetes* that includes the family of animal viruses *Astroviridae* and *Potyviridae,* the largest known family of plant viruses. We identified more than 100 family-level groups of viruses most of which have small, astrovirus-like genomes, likely, resembling the common ancestor of *Stelpaviricetes*. In contrast, the putative common ancestor of *Potyviridae* and its sister family *Potyliviridae* exhibits higher genome complexity that apparently emerged in viruses of fungi. The most plausible scenario for the origin of *Potyviridae* involves cross-kingdom horizontal virus transfer between plant-associated fungi and plants.

## INTRODUCTION

The last decade in virology was marked by an enormous expansion of the known virus diversity (1–5), a dramatic shift that occurred thanks, primarily, to the advent of virus metagenomics and metatranscriptomics (6–8). In addition to revealing a previously unsuspected vastness of the global virome, discovery of numerous new viruses in diverse environments transformed our understanding of virus ecology and informed development of fundamental concepts of the origins and evolution of viruses, and their contributions to the major transitions in the evolution of life (9–12). No less important was the application of the much improved knowledge of virus diversity to the advancement of virus megataxonomy that transformed virology from a patchwork of virus families and orders to a coherent framework based on evolutionary relationships (13). Upon acceptance by the International Committee on Taxonomy of Viruses (ICTV), this megataxonomy became the law of the land in the entire discipline (14).

Because, in a sharp contrast to the cellular organisms, there is not a single gene conserved in all viruses, viruses are obviously polyphyletic (that is, have multiple origins) precluding the construction of a single phylogenetic tree reflecting the evolution of the entire virosphere. However, expansive assemblages of diverse viruses share at least one or a few hallmark genes that can be used to generate phylogenetic trees for these broad groups. The phylogenetic tree of the hallmark gene unifying non-reverse-transcribing viruses with RNA genomes, the RNA-directed RNA polymerase (RdRP), provided a scaffold for reconstructing the evolutionary history of the RNA virosphere (15). Likewise, an RdRP homolog, reverse transcriptase (RT), was used to derive the phylogenetic tree for the RT-encoding RNA and DNA viruses (16). Taken together, these analyses resulted in the establishment of the realm *Riboviria*, which is subdivided into kingdoms *Orthornavirae* and *Pararnavirae*. The kingdom *Orthornavirae* currently consists of five large phyla further subdivided into classes, orders and families, and two more recently recognized small phyla (17).

Perhaps, the most notable novelty with respect to the composition of the global RNA virome was the discovery of a previously untapped diversity of RNA bacteriophages that now comprise about a third of all known RNA viruses (3) and feature a distinct lineage that appears to be intermediate between ‘genuine’ RNA viruses (*Orthornavirae*) and reverse transcribing viruses (*Pararnavirae*) (18). Another remarkable discovery in the RNA virosphere was the identification of the largest known RNA genomes of animal nidoviruses (19).

No less striking was the realization that the very outline of the virosphere is more nebulous than previously perceived. Apart from *bona fide* viruses defined as replicators that encode proteins encapsidating their own genomes (20), a rapidly growing number of derived viruses that encode the hallmark RdRP were found to not form virions (20). Furthermore, in addition to a handful of plant viroids that typically are minimal, non-coding circular RNA replicons, a vast variety of viroid-like RNAs was discovered, some of which encode virus capsid proteins, RdRP or uncharacterized proteins (21, 22). Thus, the previously clear boundary between non-coding viroids and capsid-encoding viruses became ambiguous, further contributing to the realization that the virosphere at large exhibits ever growing connectivity within the replicator space that includes all mobile genetic elements (20).

Understandably, preoccupation with the large-scale organization and evolution of the virosphere ran ahead of more detailed analysis of lower virus taxa such as classes and orders. In this work, we attempt at filling this gap for the ribovirian class *Stelpaviricetes* (phylum *Pisuviricota*) that includes some of the most common and economically important plant viruses. When originally established, this class consisted of two ICTV-recognized orders, *Stellavirales*, with the single familiy *Astroviridae,* and *Patatavirales*, also with a single family, *Potyviridae*. The enteric astroviruses infect a broad range of birds and mammals, and are the most common cause of diarrhea in children (23, 24). The potyviruses are the largest and most agriculturally important family of plant ribovirians (25, 26). Despite the well-established phylogenetic affinity between the astrovirus and potyvirus RdRPs (27), these two families are rather odd bed fellows. Astroviruses form icosahedral virions assembled from single jelly-roll (SJR) capsid proteins (CPs) typical of phylum *Pisuviricota* (28) and, in addition to CPs and RdRP, encode only one conserved domain, a serine protease (23, 24). In contrast, potyviruses possess a distinct type of CP that forms flexuous filamentous virions (CPf; (29, 30)) and also encode a superfamily 2 helicase (S2H) as well as two additional proteases with functions in virus replication, RNAi suppression and insect vector transmission (25, 26).

More recently, a third order, *Hypofuvirales*, with three families (*Hypoviridae, Parahypoviridae* and *Fusoviridae*) has been included in class *Stelpaviricetes* (31). This new order consists of derived, capsid-less viruses that produce both double-stranded and single-stranded forms of the genome RNA, and are associated with phytopathogenic fungi. The relationship between the first sequenced genomes of hypoviruses and potyviruses has been demonstrated in early work (32), but given the distinct biology of these viruses and the lack of capsids, they have remained formally unclassified until recently. In addition, a new family *Potyliviridae* consisting of poty-like viruses discovered by metatranscriptome mining was included in the order *Patatavirales* (31).

Here, we addressed key questions on the evolution of *Stelpaviricetes*: i) the nature of the common ancestor of this class and the ancestors of each order within it; ii) the host ranges of these ancestral viruses; iii) the evolutionary routes that led to the emergence of *Stellavirales, Patatvirales*, and *Hypofuvirales*, with their contrasting genome and virion architectures and host ranges. To address these questions, we developed a computational pipeline for comparative-genomic and phylogenetic analysis of *Stelpaviricetes*-like viruses present in the quickly growing public databases. We show that, far from being limited to 6 families, this class includes more than 100 family-level virus operational taxonomy units (vOTUs) and propose an evolutionary scenario for *Stelpaviricetes*.

## RESULTS

### *Stelpaviricetes* contigs in public databases

To represent the known diversity of viruses in the class *Stelpaviricetes*, more than 400,000 RNA virus contigs were collected from 6 datasets: ICTV, GenBank, and four datasets from extensive metagenomic surveys of ribovirians (3, 5, 33–36). To identify contigs representing *Stelpaviricetes,* the sequences were translated in six frames and compared to 121 order-specific RdRP core profiles from the Neri *et al.* study (3). The comparison was conducted using HMMER in the ‘glocal’ mode providing for the identification of complete RdRP cores (see Methods). Each contig was assigned to an order based on the hit with the highest bit-score. When this approach was applied to the kingdom *Orthornavirae* genomes from the ICTV Master Species List (MSL) 41.1 (https://ictv.global/msl) (33), RdRP was detected in 99.0% of the coding-complete genomes, and 99.1% coding-complete genomes belonging to the orders represented in the Neri *et al.* study (3) were assigned to the correct orders. Specifically, all 20 ICTV genomes from the order *Stellavirales* and 233 of the 234 ICTV genomes from the order *Patatavirales* were assigned to the correct order, whereas the 127 ICTV genomes from the order *Hypofuvirales*, which was not established at the time of the Neri *et al.* study, were assigned to orders “o.0012”, “o.0013” or “o.0014”, described by Neri *et al.* as belonging to a “base-Stelpa” class (Table S1).

Using this approach, 17,272 contigs were assigned to orders described by Neri *et al.* as belonging to the class *Stelpaviricetes* or one of the six “base-Stelpa” classes. These contigs were retained for further consideration as *Stelpaviricetes*, broadly defined (Table S2). The contigs were clustered into 3,656 species-level vOTUs based on 80% coverage and 90% identity thresholds applied to the amino acid sequences of the RdRPs (Fig. S1). A phylogenetic tree reconstructed based on RdRP domains of contigs representing the species-level vOTUs was split into family-level clades at depths locally defined by the depths of the clades encompassing the six *Stelpaviricetes* families recognized by ICTV as described in (3), yielding 103 family-level vOTUs (Fig. S1). The contribution from the Hou *et al.* (5) dataset was the largest, with its contigs represented in 93 family-level vOTUs (Fig. S2). Accumulation curves constructed by randomly sampling contigs from the dataset seem to be approaching saturation at the family level, with the total number of families extrapolated to 110, but not at the species level (Fig. S3). Although this estimate is only a rough approximation, it suggests that the substantial majority of the *Stelpaviricetes* families is already known if only as the metatrascriptome sequences.

For each vOTU, the virus with the longest RdRP-encoding contig was selected to represent that vOTU in all subsequent analyses. Notably, only 6.6% contigs representing species-level vOTUs contained a poly(A) tail. Given that genomes of most well-characterized *Stelpaviricetes* possess poly(A) tails (23, 25, 37–39), the lack of poly(A) might indicate that many if not most of the contigs do not represent complete genomes. However, it has been shown that the genomes in the highly divergent genus *Celavirus* within *Potyviridae* lack poly(A) (40), leaving uncertain the completeness of the uncharacterized genomes lacking poly(A).

This initial analysis clearly demonstrated that the evolutionary and taxonomic diversity of *Stelpaviricetes* at the family level is at least an order of magnitude greater than previously appreciated, prompting further enquiry into genome architectures, phylogeny and evolutionary history of this now vast virus class.

### Phylogenomic analysis of *Stelpaviricetes*

The RdRP-based phylogenetic tree of *Stelpaviricetes* rooted with a selection of distantly related RdRPs from the order *Picornavirales* used as the outgroup encompassed two large clades (Fig. 1). One of these clades (hereafter Astro-clade) included order *Stellavirales* (family *Astroviridae*) and a subclade formed by order *Hypofuvirales* (families *Hypoviridae*, *Parahypoviridae* and *Fusariviridae*). Another large clade (hereafter Poty-clade) included order *Patatavirales* (families *Potyviridae* and *Potyliviridae*). Of the 11 “plastroviruses” recently identified in plant transcriptomes (35) and currently unclassified, one belonged to a family-level vOTU represented by ND_179863 within the Astro-clade, whereas the rest belonged to two family-level vOTUs represented by SG001_3659 and SG001_4910, respectively, within the Poty-clade. The latter family-level vOTU, positioned at the base of the Poty-clade, also included the 42 “protopotyviruses” identified in an East China Sea aquatic RNA metavirome (41), supporting their monophyly and the apparent ancestral relationship with the potyviruses (Fig. 1).

**Fig. 1.**
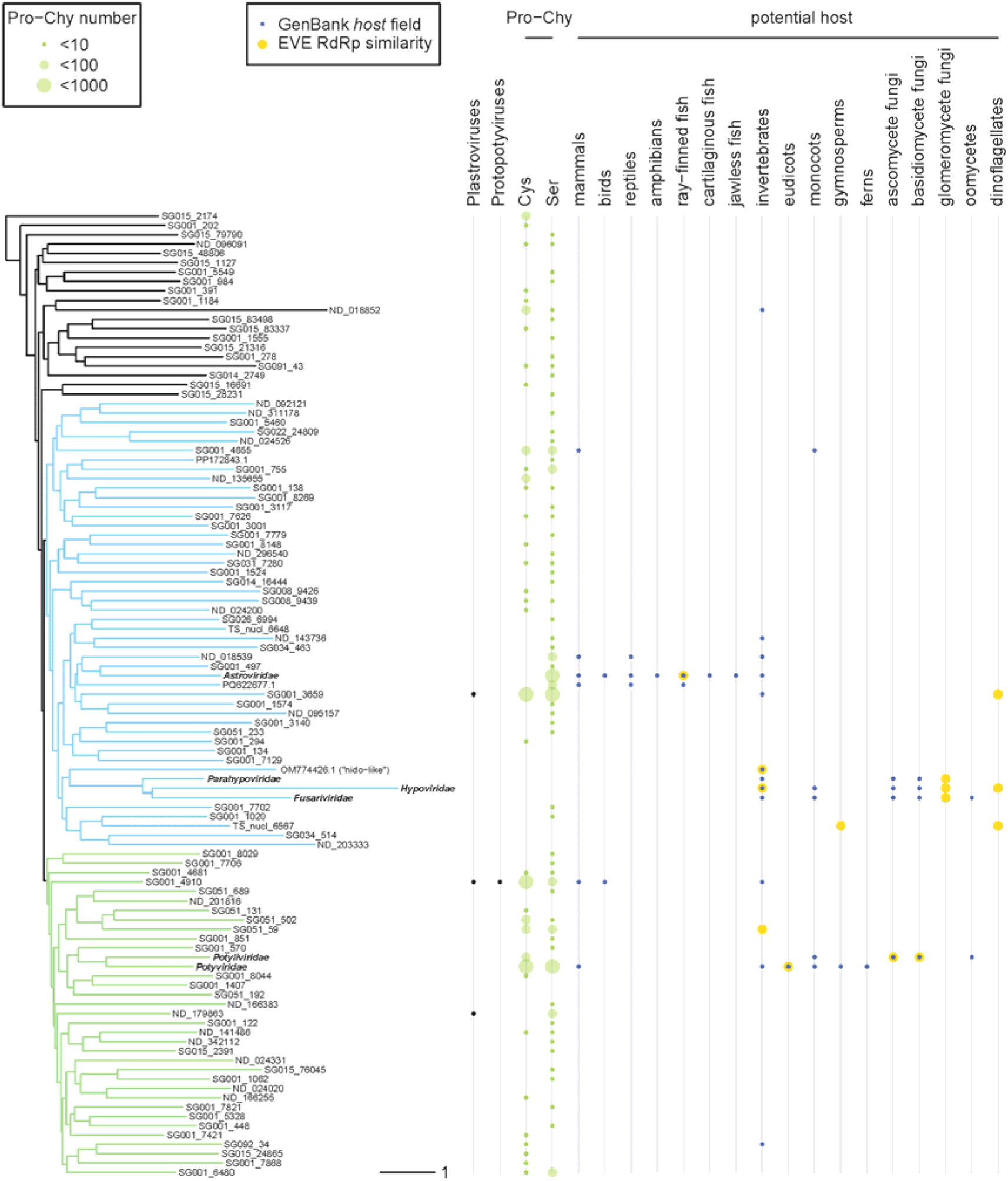
RdRP-based phylogenetic tree of *Stelpaviricetes*. One representative per family-level vOTU is shown; *Picornavirales* outgroup is not shown. Astro-clade and Poty-clade are shown in blue and in green, respectively. Hosts of vOTU members specified in GenBank are indicated by blue dots. Potential hosts of vOTU members predicted based on EVE RdRP similarity are indicated by yellow dots. Black dots indicate if any vOTU members are plastroviruses or protopotyviruses. Green ballons indicate the number of chymotrypsin-like proteases with cysteine and serine catalytic residue detected in each family-level vOTU.

Importantly, the evolutionary distances separating *Stelpaviricetes* family-level vOTUs are very large, resulting in low confidence in the reconstructed evolutionary relationships (Fig. 1). Thus, we employed tree topology testing to evaluate three alternative evolutionary scenarios: 1) common ancestry of the orders *Stellavirales* and *Patatavirales*, to the exclusion of *Hypofuvirales*, 2) common ancestry of *Stellavirales* and *Hypofuvirales*, to the exclusion of *Patatavirales,* and common ancestry of *Patatavirales* and *Hypofuvirales*, to the exclusion of *Stellavirales*. The testing methodology implemented in IQ-TREE was applied to an unconstrained RdRP-based phylogenetic tree, as well as to trees with topology constrained according to these three scenarios (Fig. S4; see Methods) (42–46). None of the scenarios could be rejected based on the test results (Table S3), implying that each one remains a possibility.

Next, we predicted the open reading frames (ORFs) and annotated protein domains encoded in the genomes representing species-level vOTUs of the expanded *Stelpaviricetes*. The genome organization of the majority of contigs was found to resemble that of *Astroviridae*, containing a 5’-terminal ORF1a encoding a chymotrypsin-like protease (Pro-Chy) domain and a downstream ORF1b encoding the RdRP that is likely expressed via a -1 ribosomal frameshift. Analogous to astroviruses, the 3’-terminal ORF2 encoding SJR CP in these contigs is most likely expressed from a subgenomic RNA (Fig. 2, Material S1-S5). In many genomes, Pro-Chy and SJR CP were not detected (Material S1), probably, due to high protein sequence divergence. Pro-Chy with either cysteine or serine catalytic residue was detected throughout the *Stelpaviricetes* tree (Fig. 1 and see below). In several instances, ORF1a and ORF1b, ORF1b and ORF2, or all three ORFs were fused into a single ORF (Fig. 2B). These variations in ORF organization were scattered throughout the RdRP-based family-level phylogenetic tree (Material S1) and even within family-level vOTUs (Material S5), suggestive of independent fusion and/or fission events occurring on multiple occasions in the evolution of *Stelpaviricetes*.

**Fig. 2.**
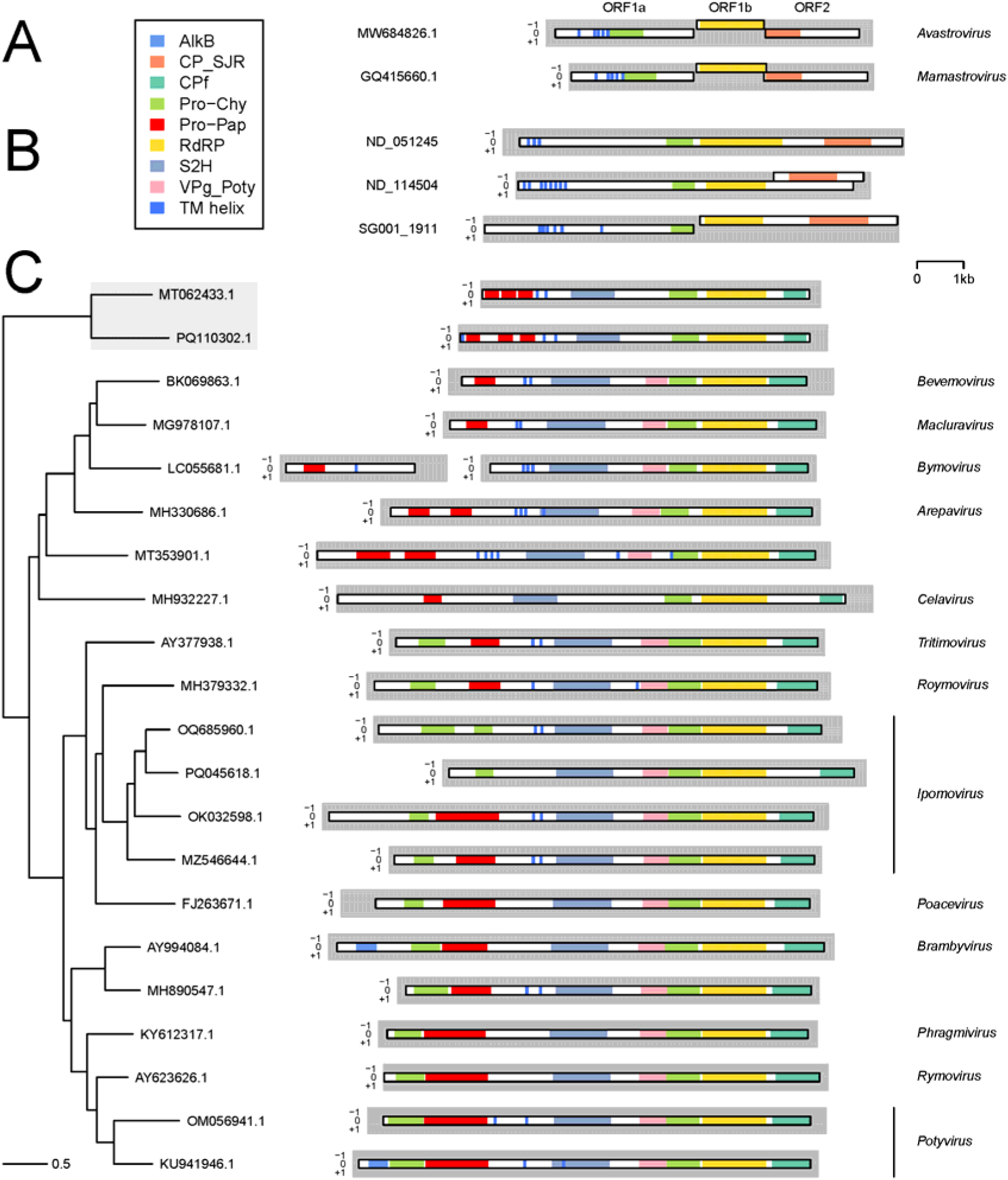
Examples of *Stelpaviricetes* genome organization. Genomes of (A) the family *Astroviridae*, (B) viruses outside of the recognized families and (C) the families *Potyviridae* and *Potyliviridae* are presented. The tree on panel C is RdRP-based, the *Potyliviridae* clade is highlighted in grey. Genera names are indicated on the right of contigs representing species-level vOTUs that include viruses classified by ICTV. Contigs are represented by grey rectangles, -1, 0 and +1 frames are shown. Frame of the most 5’-terminal ORF is set as 0. ORFs are represented by white rectangles. Protein domain annotations are indicated by color: AlkB, demethylase AlkB homolog; CP_SJR, single jelly-roll capsid protein; CPf, filamentous capsid protein; Pro−Chy, chymotrypsin-like protease; Pro−Pap, papain-like protease; RdRP, RNA-dependent RNA polymerase; S2H, superfamily 2 helicase; VPg_Poty, viral protein genome-linked of potyviruses; TM helix, transmembrane helix.

The viruses that deviated substantially from the astrovirus-like genome architecture, were the ones in vOTUs corresponding to families *Potyviridae* and *Potyliviridae* (Fig. 2C), *Hypoviridae*, *Parahypoviridae* and *Fusariviridae* (Fig. 3), the family-level vOTU represented by OM774426.1, and the ND_018852 genome (Material S1 and S5). Viruses in the *Potyviridae* and *Potyliviridae* vOTUs retain the Pro-Chy-RdRP tandem, and additionally, encode superfamily 2 helicase (S2H) upstream of Pro-Chy. Furthermore, the viruses in these two vOTUs encode CPf rather than SJR CP that is present in the vast majority of *Stelpaviricetes*. All these protein domains are encoded as a single ORF in both *Poty-* and *Potyliviridae*, further supporting the monophyly of this genome architecture.

**Fig. 3.**
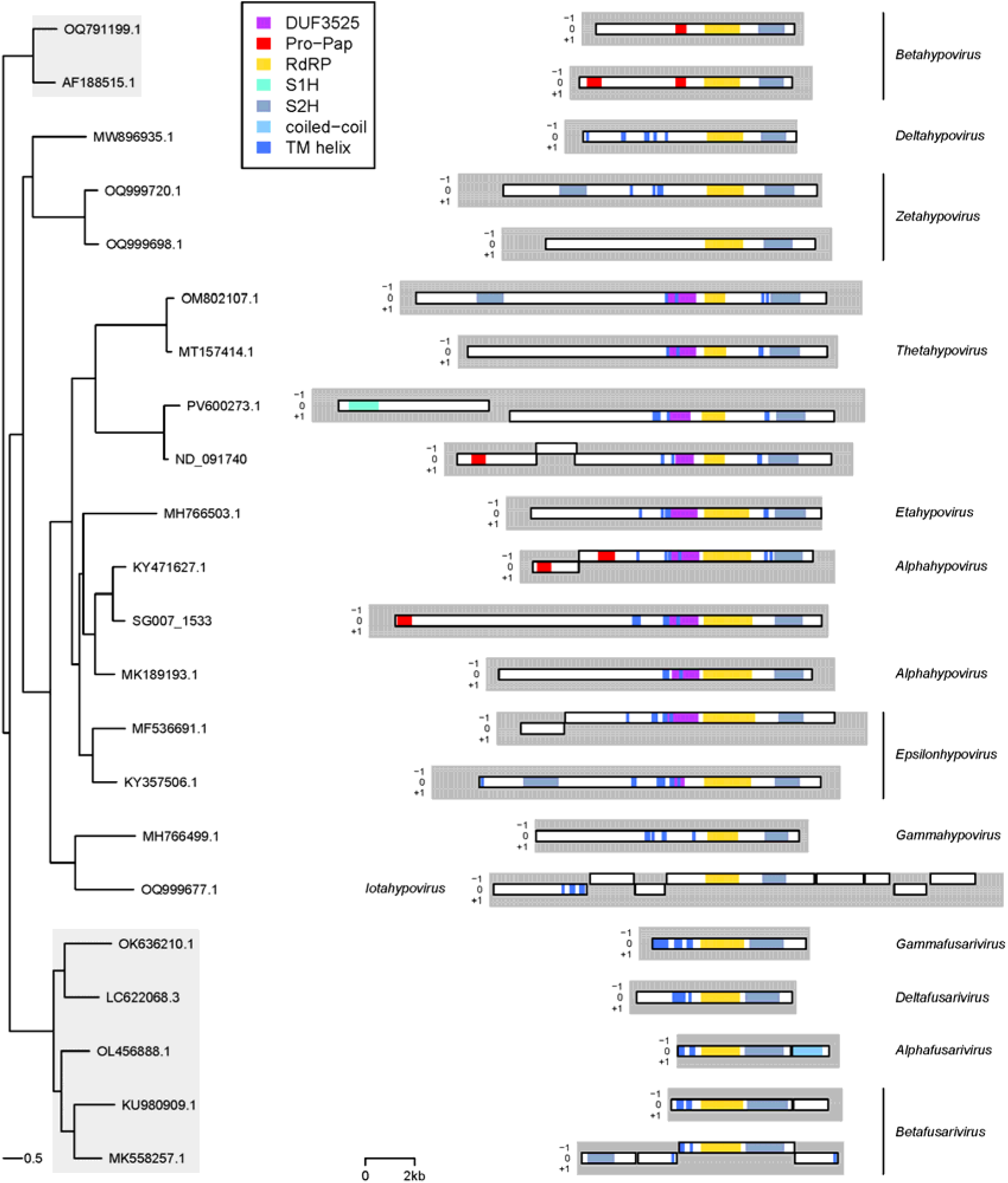
Phylogeny of *Hypofuvirales* and examples of *Hypo-, Parahypo- and Fusariviridae* genome organizations. The tree is RdRP-based, the *Parahypoviridae* and *Fusariviridae* clades are highlighted in grey, while the *Hypoviridae* clade is shown on white background. See legend of Fig. 2 for details. DUF3525, domain of unknown function; S1H, superfamily 1 helicase.

Viruses in the *Hypoviridae*, *Parahypo*viridae and *Fusariviridae* vOTUs all possess a polyprotein with an RdRP-S2H domain tandem, mimicking that of *Potyviridae* and *Potyliviridae*, albeit with the order of the domains switched. However, no capsid protein was detected in *Hypofuvirales*, and considerable variation was observed with respect to genome length, the number of ORFs and presence of additional domains (Fig. 3 and see below).

Notably, six viruses constituting the OM774426.1 vOTU possess genome organization resembling that of *Nidovirales* in *Pisuviricota* phylum, with RdRP and S1H encoded in a central ORF (Material S1, S5), and are characterized as “nido-like” in GenBank (47–51). We found that three of these viruses (OM774426.1, ND_019473 and ND_018852) additionally encode a Pro-Chy. Although they grouped with the *Hypofuvirales* clade in the phylogenetic tree, the Shimodaira-Hasegawa test support value of the grouping was low (Material S1), leaving their origin uncertain. Genome ND_018852, encoding RdRP and S1H in a single long ORF, was selected to represent a family-level vOTU as the longest genome in that vOTU, although it deviates from the astrovirus-like genome architecture of other viruses in that vOTU (Material S5). Interestingly, ND_018852 corresponds to Hubei tetragnatha maxillosa virus 7, which together with two viruses belonging to the OM774426.1 vOTU formed a clade basal to all *Nidovirales* in a previous analysis (47). The provenance of this unusual group of *Stelpaviricetes* remains to be further investigated once more genomes are sequenced.

### The host range of *Stelpaviricetes*

The hosts that are currently assigned to *Stelpaviricetes* contigs in GenBank are primarily animals, plants, fungi and oomycetes (Fig. 1). The 1,539 virus contigs identified in animal hosts belonged to 14 family-level vOTUs, with the overwhelming majority (1,464 contigs) in family *Astroviridae*. The 4,890 contigs assigned to plant hosts belonged to five family-level vOTUs, with the vast majority (4,886 contigs) in family *Potyviridae*. The 221 contigs assigned to the fungal or oomycete hosts all belonged to *Hypo-*, *Parahypo-*, *Fusari-* and *Potyliviridae* vOTUs (Fig. 1). However, the majority of the *Stelpaviricetes* contigs analyzed here originated from the metatranscriptomes and thus their hosts could not be confidently identified.

Identification of the endogenous virus elements (EVEs) integrated into genomes of cellular organisms can offer clues on the host range of related exogenous viruses. Our search across 50,401 eukaryotic datasets in the NCBI Whole Genome Shotgun database with lineage-specific RdRP profiles (3) used as queries identified 189 putative *Stelpaviricetes* EVEs. We compared the RdRP-like regions of these EVEs to the 3,656 *Stelpaviricetes* RdRP cores representing species-level vOTUs using BLASTP. As a result, 95 EVEs yielded high confidence hits (E-value <0.05, coverage of the RdRP core of a vOTU representative >100 aa). Among these hits to an EVE, the hit with the highest bit-score was used to link the EVE to an exogenous virus. Six of these 95 EVEs were very closely related to the linked viruses and consequently were excluded from consideration as likely cases of exogenous virus contamination (see Methods). The remaining 89 EVEs originated from 37 host species, including 11 animal, 3 plant, 15 fungal and 8 dinoflagellate species, and were linked to 32 *Stelpaviricetes* species-level vOTUs belonging to 10 family-level vOTUs (Fig. 1, Table S4). Notably, the host assignments for 6 of these family-level vOTUs were supported by both EVE and GenBank data (Fig. 1).

### Close relatives of *Astroviridae* in diverse vertebrates

To gain further insight into the evolutionary history of astroviruses, we analyzed the sub-tree of the RdRP-based phylogenetic tree encompassing the *Astroviridae* family-level vOTU (Fig. 4, Material S2). The sub-tree included five well-separated clades: i) the deepest-rooted clade consisting of unclassified viruses whose hosts are unknown; ii) the second deepest-rooted clade consisting of unclassified viruses hosted primarily by cartilaginous fish, amphibians and reptiles; iii) a clade consisting mostly of unclassified viruses of jawless fish and ray-finned fish; iv) clade including *Mamastrovirus* genus hosted almost exclusively by mammals; v) clade including *Avastrovirus* genus of bird viruses and unclassified viruses of amphibians, reptiles and birds (Fig. 4).

**Fig. 4.**
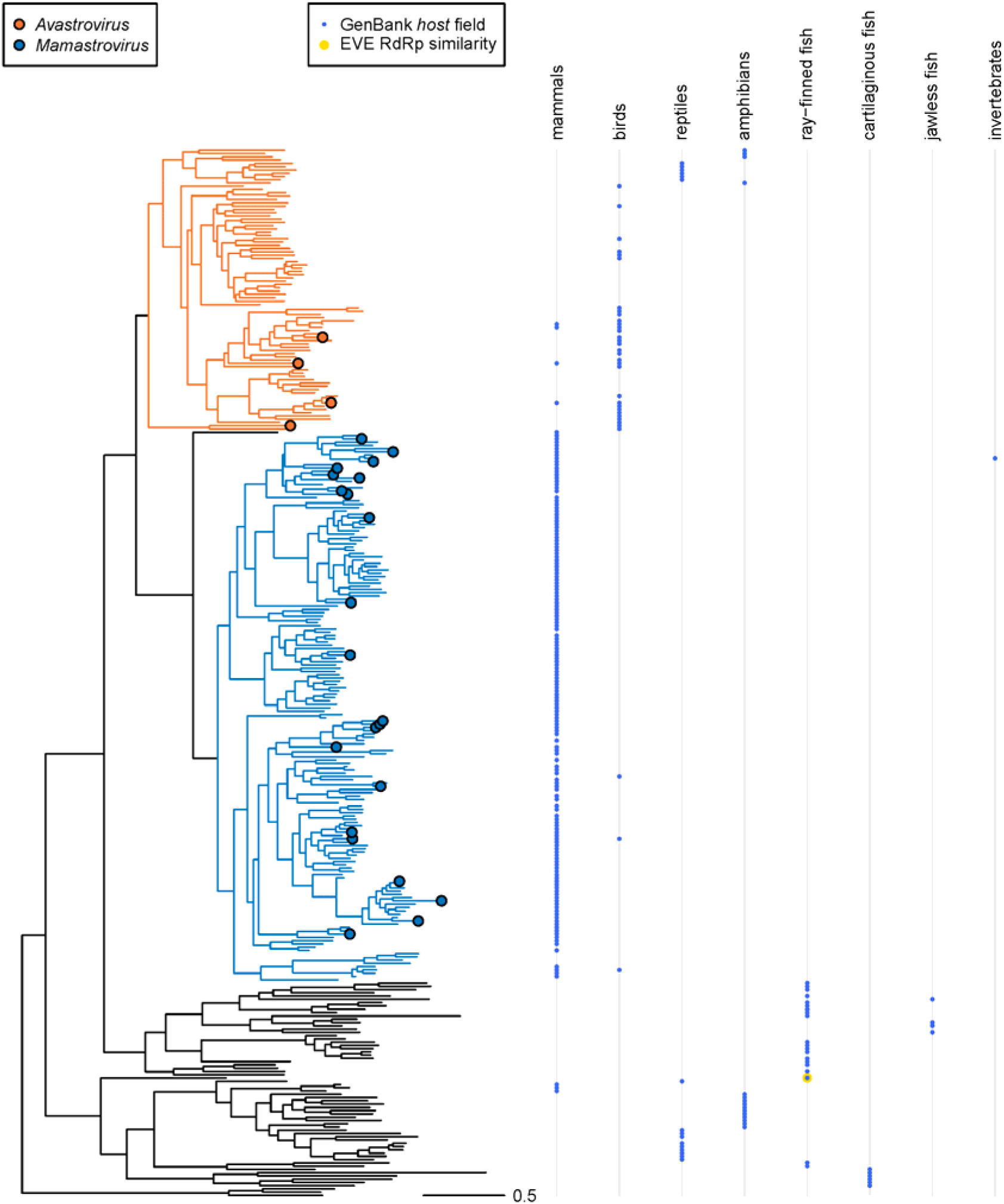
RdRP-based phylogenetic tree for *Astroviridae*. One representative per species-level vOTU is shown. The species-level vOTUs that include viruses classified by ICTV are indicated by dots colored according to ICTV genus. Corresponding clades are indicated by branch color up to the most recent common ancestor of these vOTUs. Hosts of vOTU members specified in GenBank are indicated by blue dots. Potential hosts of vOTU members predicted based on EVE RdRP similarity are indicated by yellow dots.

The topology of the tree of astroviruses and their relatives largely showed separation of the host-specific lineages, implying that these viruses co-evolved with their vertebrate hosts with limited host switching. This outcome is in agreement with the previously reported dominance of virus-host co-evolution among vertebrate ribovirians (52).

The genome architectures of the viruses in these astrovirus-like lineages mostly matched that of bona fide astroviruses, with separate ORF1a, ORF1b and ORF2 regions encoding Pro-Chy, RdRP and SJR CP, respectively (Fig. 2A, Material S2). According to an MSA of all Pro-Chy domains annotated in this study, the Pro-Chy domains throughout this family-level vOTU had serine as a catalytic residue. ORF fusion events were identified in less than 5% of the genomes representing the 320 species-level vOTUs belonging to this family-level vOTU. On the whole, the analysis of the evolution of the astroviruses shows not only coevolution with the vertebrate hosts, but also very limited innovation during the span of hundreds of millions of years.

### Origin of *Patatavirales* from fungal viruses

To explore the evolution of *Potyviridae* and *Potyliviridae*, which together constitute the order *Patatavirales*, we analyzed the sub-tree of the RdRP-based phylogenetic tree encompassing the corresponding family-level vOTUs (Fig. 5, Material S3). This sub-tree included well-separated clades matching each of the 14 ICTV-recognized potyvirus and potylivirus genera and, additionally, multiple species-level vOTUs outside of these genera (Fig. 5), further increasing the taxonomic complexity of *Potyviridae* and *Potyliviridae*.

**Fig. 5.**
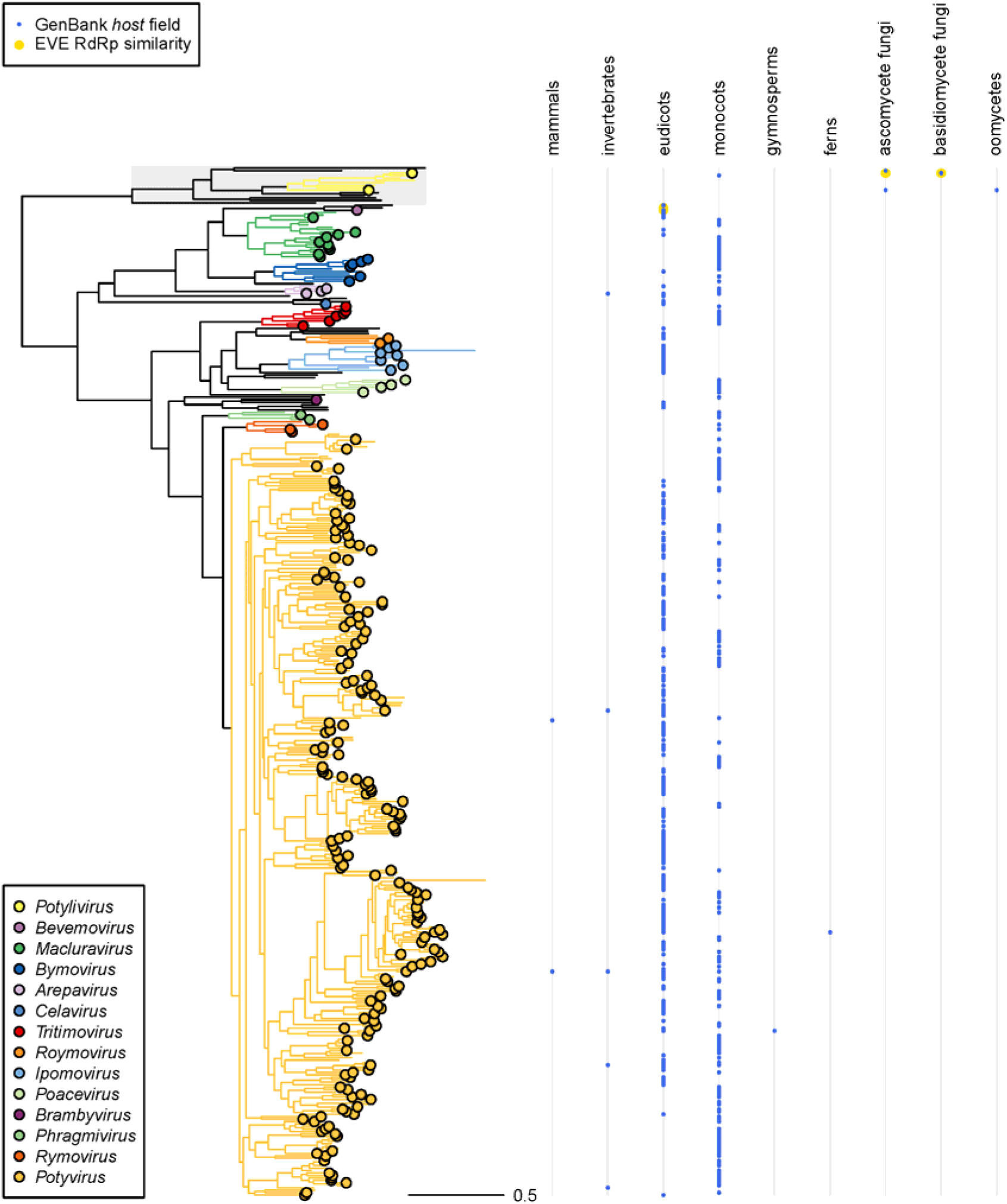
RdRP-based phylogenetic tree for *Potyviridae* and *Potyliviridae*. The *Potyliviridae* clade is highlighted in grey. One representative per species-level vOTU is shown. The species-level vOTUs that include viruses classified by ICTV are indicated by dots colored according to ICTV genus. Corresponding clades are indicated by branch color up to the most recent common ancestor of these vOTUs. Hosts of vOTU members specified in GenBank are indicated by blue dots. Potential hosts of vOTU members predicted based on EVE RdRP similarity are indicated by yellow dots.

The genomes of both *Potyviridae* and *Potyliviridae* encompass a single long ORF, with the exception of the genus *Bymovirus* in which this ORF is split between two genome segments. In addition to the Pro-Chy-RdRP tandem that is conserved in most *Stelpaviricetes,* all complete virus genomes in this sub-tree encoded S2H and CPf (Fig. 2C; Material S3). The Pro-Chy domains that are present in all *Potyviridae* and *Potyliviridae* vOTUs upstream of the RdRP invariably contained cysteine as the catalytic residue.

The organization of the 5’-terminal region of the viral genomes within *Patatavirales* varies with respect to the number and type of encoded proteases. Almost all members of the family *Potyviridae* encode, in the N-terminal region of the polyprotein, one or two papain-like cysteine proteases (Pro-Pap) known as helper component-proteinases (HC-Pro) (53). In one of the two major lineages of *Potyviridae*, including the genus *Potyvirus*, HC-Pro is preceded by another, diverged chymotrypsin-like serine protease known as P1 (54, 55). Some of the *Ipomovirus* genus members lack HC-Pro, and a subset of these encodes a third chymotrypsin-like serine protease in the polyprotein N-terminus (Fig. 2C; Material S3) (56). Examination of the MSA of all Pro-Chy domains annotated in this study, demonstrated that in all but one potyvirus N-terminal Pro-Chy, the catalytic residue is serine.

The information on the domain organization of the polyprotein N-termini in the genus *Celavirus* and the family *Potyliviridae* is scarce. Furthermore, the comparison of the polyprotein sequences from these groups with the viral protein profile database NVPC [9] did not yield reliable functional annotations of the polyprotein N-termini, indicative of substantial divergence in this region. To investigate the domain organization of these polyprotein N-termini, we extracted N-terminal regions of the polyproteins upstream of S2H from viruses representing species-level vOTUs, namely, three celaviruses and 11 potyliviruses. The structures of these regions were predicted using AlphaFold3 and the resulting structural models were compared to the Protein Data Bank (PDB) protein structure database (57). All celaviruses showed significant structural similarity to the turnip mosaic virus (TuMV) HC-Pro (PDB 3RNV; Table S5) [10]. The catalytic core of the putative proteases of celaviruses could be confidently superimposed with the 3RNV structure (Fig. S5) and was conserved in the alignment of the N-terminal regions of the celavirus polyproteins (Fig. S6A, Table S5).

Notably, in the N-terminal regions of the polyproteins of potyliviruses, we identified three consecutive papain-like protease domains (Fig. 2C). The first, second and third predicted proteases were detected through significant structural similarity to the TuMV HC-Pro structure in 5, 9 and 7 potyliviruses, respectively (Table S5), and the catalytic cores of these predicted proteases could be superimposed with the 3RNV structure in all but one case (Table S5). The three papain-like proteases were conserved in the alignment of the N-terminal regions of the potylivirus polyproteins (Fig. S6B-D), and the alignment conservation allowed us to extend the first and third protease domain detection to one and two more viruses, respectively (Fig. S6B and S6D, Table S5). While the three proteases were detected in six of the 11 analyzed potyliviruses, in three other viruses, only the second and third protease were detected, and in two viruses, no papain-like proteases were identified; we assume that these differences were due to the incompleteness of the respective virus genomes at the 5’-terminus (Material S3). Thus, the vast majority of the viruses in the families *Potyviridae* and *Potyliviridae* harbored at least one homolog of the potyvirus HC-Pro, which is duplicated in the *Arepavirus* genus and in one of the viruses outside of the recognized genera of *Potyviridae* (MT353901.1; Fig. 2C), and triplicated in *Potyliviridae*.

The 30 genomes-strong *Potyliviridae* family included five viral genomes derived from metatranscriptomes enriched in fungal and oomycete sequences (Fig. 5, Table S2) (58–61). Furthermore, EVEs closely related to one of the viruses in this lineage (MK231047.1) were identified in genomes of six fungal species that belong mostly to parasitic ascomycetes of the *Tolypocladium* genus (Fig. 5, Table S4). One of the viruses in this clade, Macrophomina phaseolina poty-like virus (MT062433.1), was isolated from a cultured plant-pathogenic ascomycete *Macrophomina phaseolina*, providing further evidence for fungi being the actual hosts of the viruses in this family (60), and by extension, a likely fungal origin of *Patatavirales*.

The phylogenomic distribution of protein domains encoded by *Potyviridae* and *Potyliviridae* analyzed above (Fig. 2C; Material S3) implies monophyly of HC-Pro, S2H and CPf that, most likely, were acquired by the common ancestor of *Patatavirales*. By contrast, P1 likely emerged in the common ancestor of one of the two major lineages within *Potyviridae* (Fig. 2C; Material S3).

### Phylogenomics of *Hypofuvirales*

To explore the evolution of the viruses in the order *Hypofuvirales*, we analyzed the sub-tree of the RdRP-based phylogenetic tree encompassing the corresponding family-level vOTUs (Fig. 6, Material S4. Family *Parahypoviridae* vOTU was basal to a clade formed by *Hypo-* and *Fusariviridae* family-level vOTUs. The genera of the three families formed well-separated clades, with the exception of the genus *Iotahypovirus* of the family *Hypoviridae*, the members of which were found to be scattered across the *Hypoviridae* family clade. Besides, there were multiple species-level vOTUs outside of the genera recognized by ICTV.

**Fig. 6.**
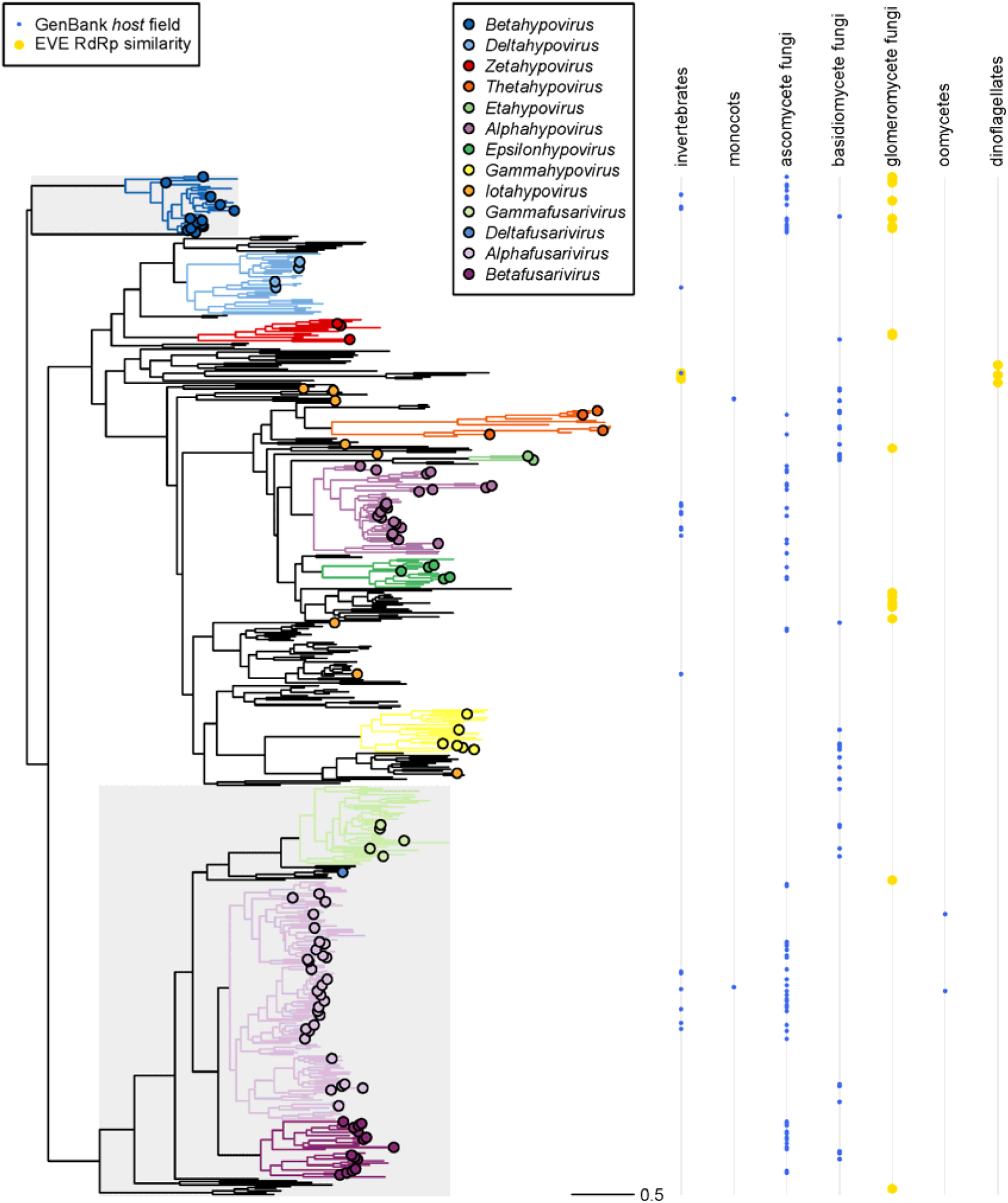
**RdRP-based phylogenetic tree for *Hypoviridae, Parahypoviridae and Fusariviridae***. The *Parahypoviridae* and *Fusariviridae* clades are highlighted in grey, while the *Hypoviridae* clade is shown on white background. One representative per species-level vOTU is shown. The species-level vOTUs that include viruses classified by ICTV are indicated by dots colored according to ICTV genus. Corresponding clades are indicated by branch color up to the most recent common ancestor of these vOTUs. Note that the genus *Iotahypovirus* is non-monophyletic. Hosts of vOTU members specified in GenBank are indicated by blue dots. Potential hosts of vOTU members predicted based on EVE RdRP similarity are indicated by yellow dots.

GenBank metadata pointed to ascomycete and basidiomycete fungi as the most prominent hosts of *Hypofuvirales* (Fig. 6). Additionally, we identified EVEs closely related to *Hypofuvirales* in glomeromycete fungi and dinoflagellate genomes (Fig. 6; Table S4).

There is substantial variation in genome organization among the *Hypofuvirales* (Fig. 3; Material S4). All complete *Hypofuvirales* genomes representing species-level vOTUs encode a RdRP-S2H domain tandem embedded in a polyprotein, and no capsid protein domains were detected. Apart from this conserved core, the *Hypofuvirales* genomes varied in terms of length (reaching 22,4 kb), number of ORFs (reaching 8 ORFs) and presence of additional protein domains (62). Some of the viruses in the genera *Thetahypovirus* and *Alphahypovirus* of the family *Hypoviridae* harbored a single Pro-Pap in the polyprotein N-terminus, whereas the KY471627.1 hypovirus genome (63) was found to encode an additional Pro-Pap in a small 5’-terminal ORF. Notably, this domain architecture with two Pro-Pap domains was first described for the Cryphonectria hypovirus 1 genome M57938.1, which belongs to the species-level vOTU represented by KY471627.1, emphasizing the similarity of the polyprotein organization to that of potyviruses (32, 37). Furthermore, nearly all members of the *Parahypoviridae* family-level vOTU were found to encode a permuted Pro-Pap domain upstream of the RdRP (64), whereas parahypovirus genomes AF188515.1 and PP999648.1 each possessed an additional regular Pro-Pap domain positioned in the polyprotein N-terminus. In addition, several genomes in the *Hypoviridae* and *Fusariviridae* families encode a second helicase domain in the 5’-terminal region of the genome (62, 65, 66).

### Evolutionary relationships of unique *Stelpaviricetes* domains

To analyze the evolutionary history of domains found in *Stelpaviricetes* whose genomes deviate from the characteristic Pro-Chy-RdRP-SJR astrovirus-like domain architecture, we clustered sequences of *Stelpaviricetes* Pro-Pap, S1H, S2H and CPf domains and compared the representatives of the clusters to the NCBI Protein Reference Sequences database using BLASTP.

We identified thousands of S1H and S2H homologs in cellular organisms and viruses and used their sequences to reconstruct phylogenetic trees. Notably, bacterial HrpB helicases were prevalent among the identified S2H homologs. In the S1H phylogenetic tree (Fig. S7), S1H of ND_018852 and the family-level vOTU represented by OM774426.1 grouped with S1H of *Nidovirales*, whereas the two S1H domains detected in *Hypoviridae* grouped together and were extremely distant from all other S1H helicases.

In the S2H phylogenetic tree (Fig. S8), S2H domains of *Hypofuvirales* and *Patatavirales* formed two major clades. Within the *Hypofuvirales* clade, S2H domains of *Hypoviridae* and *Parahypoviridae* grouped together, whereas S2H domains of *Fusariviridae* formed a separate sub-clade, and the S2H domains encoded in the 5’-termini of several *Hypo-* and *Fusariviridae* genomes constituted a separate sub-clade as well. The S2H domains of *Patatavirales* formed a strongly supported clade that at its base included S2H domains of class *Flasuviricetes* (flavivirus-like viruses), the only other group of ribovirians encoding S2H. Curiously, S2H domains of the genus *Celavirus* clustered with those of *Flasuviricetes*. There is no obvious outgroup for the S2H phylogeny, and therefore, the tree remained unrooted. Nevertheless, the ribovirian S2H clade was separated from the cellular ones by a long branch, suggesting that the ribovirian S2H are monophyletic and spread horizontally between distant groups of viruses, such as *Patatavirales* and *Flasuviricetes*.

In contrast, no non-*Stelpaviricetes* homologs of Pro-Pap or CPf were identified using the above approach, even when PSI-BLAST with three iterations was used instead of BLASTP. The only exception was the protease domain of Sclerotinia sclerotiorum megabirnavirus 1, for which an acquisition through horizontal gene transfer had previously been suggested (67). Consequently, to identify distant homologs of these domains, we used AlphaFold3 to predict structures of the representative domain sequences and compared the predicted structures to the PDB database.

Non-permuted Pro-Pap models produced significant hits to potyvirus (3RNV-A) and arterivirus (3MTV-A) papain-like proteases, although the structures showed substantial differences (Fig. 7A-D) (68, 69). The permuted Pro-Pap model showed significant structural similarity to a murine deubiquitinase (2WP7-A; Fig. 7E-F), *Vibrio cholerae* type VI secretion system effector (6V98-A) and human ceroid lipofuscinosis neuronal protein 5 (6R99-A) (70–72). CPf models, apart from the obvious hit to the potyvirus CPf structure (8ACC-A), showed significant similarity to the alphaflexivirus CPf (5FN1-A, 5A2T-A) (30, 73, 74) and bunyavirus nucleoprotein (4CSF-K) as reported previously (30, 75–77). These findings, like the S2H analysis, suggest horizontal gene transfer between distant ribovirians as well as independent acquisition of permuted Pro-Pap from the host.

**Fig. 7.**
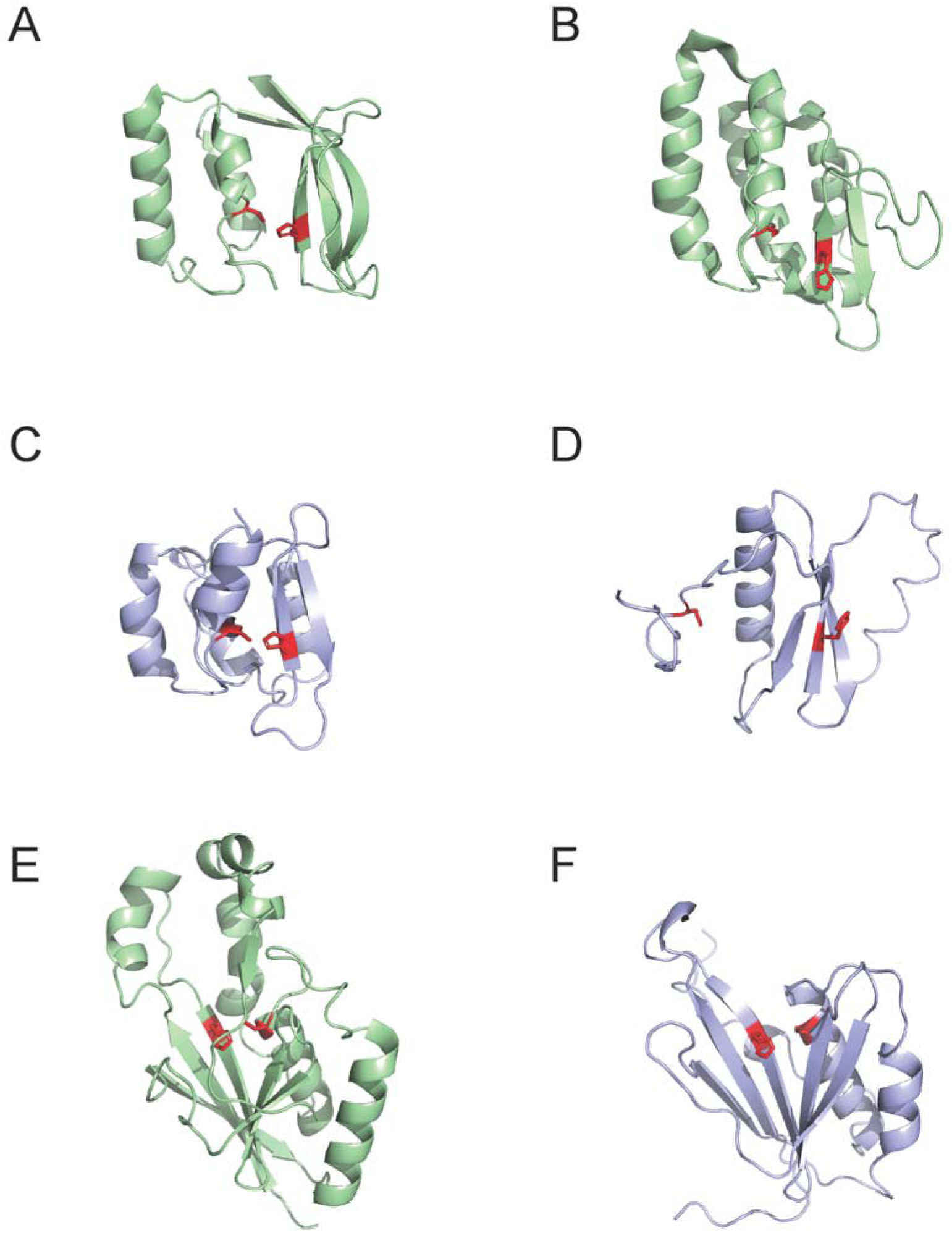
Structures of papain-like proteases. Crystal structures (green) and models (blue) of Pro-Pap of (A) porcine reproductive and respiratory syndrome virus (3MTV-A), (B) turnip mosaic virus (3RNV-A), (C) uromyces potyvirus A (MK231047.1, polyprotein fragment 287-379 aa), (D) fusarium graminearum hypovirus 1 (MK279472.1, polyprotein fragment 127-213 aa), (E) murine deubiquitinating isopeptidase (2WP7-A) and (F) fusarium oxysporum dianthi hypovirus 2 (MN176979.1, polyprotein fragment 1309-1443 aa). Catalytic dyad is highlighted in red.

## DISCUSSION

The dramatic expansion of the collection of virus genomes enabled by the advances of metagenomics and, in the case of ribovirians, metatranscriptomics provides both unprecedented opportunities and considerable challenges to the study of the diversity and evolution of the global virome (1–8). The known genomic diversity of viruses is burgeoning at all taxonomic levels, and in order to reconstruct the evolutionary events occurring within various viral clades and identify general patterns and trends, detailed phylogenomic analysis is indispensable (15). In this study, we were specifically interested in the class *Stelpaviricetes* (phylum *Pisuviricota*) for two primary reasons. First, this class includes *Potyviridae*, the largest, most diverse and abundant family of plant ribovirians of considerable agricultural importance (25) evolutionary origin of which remained obscure. Second, *Stelpaviricetes* includes viruses with distinct genome architectures, namely, relatively complex genomes of potyviruses encapsidated into filamentous virions and much smaller genomes of astroviruses encapsidated into icosahedral virions (23, 24), making this class an interesting target for evolutionary reconstruction. Thus, the major goal of this work was to investigate the much-expanded known diversity of the *Stelpaviricetes*, in an attempt to uncover its evolutionary history.

The first striking outcome of our phylogenomic analysis was that, rather than being limited to the six currently recognized families of *Astroviridae, Potyviridae, Potyliviridae, Hypoviridae, Parahypoviridae* and *Fusariviridae*, *Stelpaviricetes* now encompass more than 100 (putative) families, thus expanding the known family-level diversity of this class by roughly an order of magnitude. The accumulation curves for *Stelpaviricetes* (Fig. S3B) suggest that relatively few family-level clades remain to be discovered.

Equally notable, despite the extensive divergence of the RdRP sequences, all but five of the 103 family-level vOTUs displayed a near uniform genome architecture closely resembling that of astroviruses, with only three recognizable protein domains, RdRP, Pro-Chy and SJR CP. Among these, RdRP is universally conserved in ribovirians whereas SJR CP and Pro-Chy are among the most common domains in the collective ribovirian proteome (78). Compared to the reconstructed common ancestor of the ribovirians of eukaryotes that was inferred to have encoded only two protein domains, RdRP and SJR CP (15), a typical *Stelpaviricetes* genome has acquired a protease domain, one of the prerequisites for the further ribovirus genome complexification (11).

The only type of deviations from that common genome architecture of *Stelpaviricetes* that independently recurred in multiple clades involved ORF fusions so that these domains can be contained in three (the most widespread, likely, ancestral configuration), two or a single protein. A sharp departure from this uniform architecture was observed in two major clades of *Stelpaviricetes* corresponding to the orders *Patatavirales* and *Hypofuvirales*. Given that in the *Stelpaviricetes* tree, these clades do not emanate from the root but rather are embedded within larger lineages of viruses with the astrovirus-like genomes (Figs. 1 and 2), it appears most likely that the common ancestor of the class had this signature, astrovirus-like architecture. At the base of the *Patatavirales* clade, three domains were apparently acquired in concert, Pro-Pap, S2H and CPf, the latter replacing the ancestral SJR CP. Of these, a helicase domain, together with the proteases, was identified as another major prerequisite of genome complexification in ribovirians (11, 79). Together, such domains contribute to increased efficiency and flexibility of the RNA virus genome replication, expression and interactions with the host (11).

The evolutionary provenance of *Hypofuvirales* remains uncertain given the lack of support for the tested alternative topologies of the RdRP phylogeny. The presence of S2H in all and Pro-Pap in some members seems to suggest origin from *Patatavirales* (possibly, *Potylivirdae* with their fungal hosts) as proposed in early work (32). This evolutionary scenario would involve loss of Pro-Chy and CPf, and switch to a capsidless lifestyle. However, given the change of the order of RdRP and S2H domain, and the lack of clear affinity between the S2H and Pro-Pap of *Patatavirales* and *Hypofuvirales*, alternative, more complex evolutionary scenarios cannot be ruled out.

Elucidation of the specific origin of the novel domains captured by the ancestor of mycopotyvirus/potyvirus lineage is a challenging task given the high sequence divergence typical for ribovirus evolution (15). Remarkably, the S2H of the *Patatvirales* lineage appears to share a common ancestor with the helicases of *Amarillovirales* (phylum *Kitrinoviricota*, class *Flasuviricetes*), the only known group of ribovirians outside *Stelpaviricetes* encoding S2H, suggesting gene exchange between these viruses from distinct ribovirian phyla (17). For Pro-Pap and CPf, the results of structural comparisons were less clear, but it is notable that in both cases, the highest structural similarity was again observed with proteins from distantly related viruses (including a different phylum*, Negarnaviricota*, in the case of CPf), further emphasizing the likely role of interviral gene exchanges in the origin of major groups of viruses (11).

The difficulties in identifying the exact sources of additional domains in *Patatvirales* emergence notwithstanding, it should be emphasized that the closest relatives of these domains were identified in diverse ribovirians whose host ranges include protists, animals, fungi and plants (see the closing section of Results). Such breadth of the host range of viruses potentially contributing to the evolution of *Patatavirales* suggests extensive horizontal gene transfer among diverse ribovirians, either co-infecting the same host organisms or being in contact via close ecological associations among their respective hosts. The latter possibility is particularly relevant for fungi whose life styles routinely involve symbiotic relationships with diverse organisms, from protists to animals to plants (80–82).

For most of the viruses identified in metatranscriptomes, host assignment is unfeasible. Nevertheless, combining EVE analysis with information on isolated viruses yields some valuable clues. In particular, the *Potyliviridae*, the sister clade of *Potyviridae* is likely hosted by fungi (and possibly, oomycetes as well), suggesting that the common ancestor of *Patatavirales* replicated in fungi, and the ancestor of the *Potyviridae* later transferred to plants. This sequence of events is compatible with the well-established fact that fungi and their virome have emerged and diversified much earlier than land plants (83, 84). The subsequent co-terrestrialization of fungi and plants followed by emergence and explosive diversification of the flowering plants resulted in close ecological association between many species from these eukaryotic kingdoms. This association involves all types of plant-fungus symbioses including parasitism, commensalism and mutualism, providing ample opportunities for bidirectional horizontal virus transfer between fungi and plants, as proposed previously (85), as well as gene exchange among diverse viruses. Our phylogenomic analysis of *Stelpaviricetes* provides additional support for the concept of fungal viruses being a significant contributor to the evolution of the land flora virome, in addition to a dominant contributions from the virome of invertebrates.

Furthermore, the EVE analysis presented here suggested that some viruses with the typical astrovirus-like genome organization replicate in fungi and dinoflagellates, implying that the common ancestor of *Stelpaviricetes* might have been a protist virus. The origin of the vertebrate astroviruses is difficult to reconstruct due to the limited information on the host specificity of the sister clades of *Astroviridae*. The identification of several astrovirus-like viruses in the invertebrate transcriptomes (1, 47) seems to suggest that the ancestors of *Astroviridae* were hosted by invertebrates. In general, however, host assignment for the majority of the newly discovered ribovirians, which is essential for confident inference of evolutionary scenarios, remains elusive. This challenge has to be addressed by screening larger databases of eukaryotic genomes for EVEs and by developing much needed large-scale approaches for detecting virus-host associations. Nevertheless, the recent major expansion of the established host range of *Stelpaviricetes*, from merely birds, mammals and plants to fish, amphibians, invertebrates, fungi, oomycetes, and dinoflagellates, as shown in this and other studies (36, 86), greatly enriches the picture of the *Stelpaviricetes* ecology and evolution.

### Conclusions

In this study, we dramatically expanded the taxonomic breadth of the ribovirus class *Stelpaviricetes* at the family level, from six currently recognized families to 103. We also attempted to answer the three central questions on the evolution of *Stelpaviricetes.* First, we propose that the common ancestor of this virus class was a protist-infecting astrovirus-like virus that encoded RdRP, Pro-Chy and SJR CP domains. Second, our phylogenomic analyses suggest that the common ancestors of vertebrate *Astroviridae* and plant *Potyviridae* were hosted by invertebrates and fungi, respectively. Third, we provide substantial evidence that, in contrast to astrovirus-like viruses with their small genomes and limited domain repertoire that comprise the bulk of families in this, now vast class of ribovirians, the more complex genomes of *Patatavirales* emerged via acquisition of the Pro-Pap and S2H domains and replacement of SJR CP with CPf. These evolutionary innovations most likely occurred in the fungal ancestors of *Patatavirales*, possibly, via gene exchange with diverse viruses. Although the reliability of the presented evolutionary scenarios remains to be further probed along with the expansion of ribovirian sampling throughout eukaryotic hosts, this study provides a framework for future phylogenomic analyses of the expanded *Stelpaviricetes*.

## METHODS

### Datasets

Kingdom *Orthornavirae* genome sequences from the ICTV MSL 41.1 (n = 11,775) and GenBank (n = 203,516; >3 kb, SARS-CoV-2, hepatitis C virus and human respiratory syncytial virus genomes excluded, downloaded on 15 September 2025), plastrovirus contigs described in Lauber et al. 2019 (n = 11), Neri et al. 2022 contigs >3 kb described as belonging to the class *Stelpaviricetes* or one of the six “base-Stelpa” classes (n = 2,093), as well as RNA virus contigs >3 kb from Hou et al. 2024 (n = 190,302) and Liu et al. 2026 (n = 3,954) datasets were analyzed (3, 5, 33–35). The GenBank metadata was extracted using NCBI Entrez 24.7 and NCBI Datasets 18.18.0 (87). Sequences were processed with the help of SeqKit 0.11.0, Seqtk 1.4 (https://github.com/lh3/seqtk), R packages SeqinR 4.2-36 and Bio3D 2.4-4 (88, 89).

### RdRP core delineation and classification

Virus genome sequences were translated in six frames using EMBOSS 6.6.0 ‘transeq’ with the ‘-frame 6 -clean Y -alternative Y’ parameters (90). The translations were compared to 121 HMM profiles built based on order-specific RdRP core MSAs from Neri et al. (3). The comparisons were conducted by HMMER 2.3.2 ‘hmmsearch’ in ‘glocal’ mode aimed to provide a query-target alignment global in respect to the query HMM and local in respect to the target protein sequence (http://hmmer.org/). E-value 0.05 and bit-score 0 thresholds were applied. For each virus genome sequence, the RdRP core region was delineated and assigned to an order based on the hit characterized by the highest bit-score.

The delineated RdRP core sequences assigned to orders described by Neri et al. as belonging to the class *Stelpaviricetes* or one of the six “base-Stelpa” classes were retained for further consideration unless they (1) contained >5% ‘X’ symbols or (2) when aligned using Muscle 5.3 ‘Super5’ algorithm (91), contained a residue other than aspartate in the column corresponding to the N-terminal conserved aspartate of the RdRP motif A or C, or a residue other than aspartate or asparagine in the column corresponding to the C-terminal conserved aspartate of the RdRP motif C, or a gap in the column corresponding to the conserved threonine of the RdRP motif B (92).

### RdRP core clustering

The delineated *Stelpaviricetes* RdRP core sequences were clustered into species-level vOTUs using MMseqs2 3.0 with the ‘--cov-mode 1 -c 0.8 --min-seq-id 0.9’ parameters (93). Next, an RdRP-based phylogenetic tree of species-level vOTU representatives was split into family-level clades at depths locality-dependent on that of the clades encompassing the six *Stelpaviricetes* families recognized by ICTV, as described in (3). In each vOTU, the virus with the longest RdRP-encoding contig was selected to represent the vOTU in subsequent analyses. If the RdRP domain was encoded in a reverse frame of the original contig sequence, a reverse complement sequence was analyzed.

### RdRP-based phylogeny reconstruction

RdRP core sequences of the representatives of species-level vOTUs, as well as RdRP core sequences of poliovirus 1 Mahoney (V01149.1) and cowpea mosaic virus (X00206.1) serving as an outgroup, were aligned using Muscle 5.3 ‘Super5’ algorithm (91). Alignment columns containing >50% gaps were excluded from consideration. The MSA was inspected using Jalview 2.11.4.1 (94). A phylogenetic tree based on the MSA was reconstructed using FastTree 2.2.0 with the ‘-lg’ parameter (95). All RdRP-based trees analyzed in this project were derived from the reconstructed tree using R packages Ape 5.8 and Phytools 2.3-0 (96, 97).

### Tree topology testing

The dataset included 50 phylogenetically distant representatives from each of the orders *Stella-*, *Patata-* and *Hypofuvirales*, one representative from each of the 97 *Stelpaviricetes* family-level vOTUs outside of the three ICTV-recognized orders, as well as two order *Picornavirales* viruses, poliovirus 1 Mahoney and cowpea mosaic virus. Phylogenetically distant representatives were selected by extracting a clade corresponding to an order from the *Stelpaviricetes* RdRP-based phylogenetic tree reconstructed as described above, measuring pairwise distances between the clade members (R package Ape 5.8 function ‘cophenetic.phylo’), hierarchically clustering the clade members based on the pairwise distances (R function ‘hclust’), cutting the hierarchical clustering dendrogram to produce 50 clusters (R function ‘cutree’) and selecting one representative per cluster (96). The alignment used to reconstruct the trees was a subset of the RdRP core sequences alignment described above, columns containing >50% gaps were removed prior to tree reconstruction. A tree was reconstructed without imposing a constraint on its topology using IQ-TREE 2.1.3 with the ‘--mset WAG,LG’ parameter; the “LG+F+R9” model was selected. Topology of three trees reconstructed using the IQ-TREE with the ‘-m LG+F+R9’ parameter was constrained as follows: i) orders *Stellavirales* and *Patatavirales* belong to sister clades, *Hypofuvirales* is basal to them, *Picornavirales* is basal to these three *Stelpaviricetes* orders; ii) order *Stellavirales* and *Hypofuvirales* belong to sister clades, *Patatavirales* is basal to them, *Picornavirales* is basal to these three *Stelpaviricetes* orders; iii) orders *Patatavirales* and *Hypofuvirales* belong to sister clades, *Stellavirales* is basal to them, *Picornavirales* is basal to these three *Stelpaviricetes* orders. IQ-TREE phylogenetic testing was applied to the four trees using ‘-n 0 -zb 10000 -au’ parameters (42–46).

### Accumulation curve construction

Accumulation curves were constructed using function ‘specaccum’ (method ‘random’ and 100 permutations) from the R package Vegan 2.7-2. The family accumulation curve was approximated by the Michaelis-Menten model using R function ‘nls’, allowing to estimate the total number of families as an asymptote of the model.

### EVE comparison

Endogenous RdRP-like regions were identified by initially searching for RdRP sequences from Neri et al. (3) against the 50,401 eukaryotic datasets in the NCBI Whole Genome Shotgun database using MMseqs2 (93) and subsequently verifying the matches by running lineage-specific RdRP profiles (3) against the 6-frame translation of the respective contigs using PSI-BLAST (98). The identified 189 *Stelpaviricetes* EVE RdRP-like regions were compared to 3,656 *Stelpaviricetes* RdRP cores representing species-level vOTUs using BLASTP 2.17.0+ (99). Only the hits characterized by E-value <0.05 and covering >100 aa of a *Stelpaviricetes* RdRP core were considered. In total, 95 out of the 189 EVEs received hits meeting these criteria. For each of the 95 EVEs, the hit characterized by the highest bit-score was used to link it to a virus. Six EVE RdRP-like regions were found to be highly similar to the RdRPs of the linked viruses (>90% aa identity). The contigs harboring these six EVEs were short (<10 kb) and highly similar (>80% nt identity along the entire length of the EVE-containing contig) to the genomes of the linked viruses. Consequently, these six EVEs were excluded from consideration as likely cases of exogenous virus contamination. The source organisms of the remaining 89 EVEs were regarded as potential hosts of the linked viruses.

### Polyadenylated tail detection

A contig was considered to possess a poly(A) tail if there were >10 adenine nucleotides at the 3’-end.

### ORF prediction

ORFs of the genomes representing species-level vOTUs were predicted by EMBOSS 6.6.0 ‘getorf’ with the ‘-find 1 -table 0 -minsize 600’ parameters (90).

### Transmembrane regions prediction

Transmembrane helices in the ORF products were predicted by TMHMM 2.0 (100).

### Sequence-based protein domain annotation

A database composed of 4,833 NVPC database profiles (3) and 75 Pfam 38.2 database profiles (30 profiles from the Pro-Chy clan CL0124 and 44 profiles from the Pro-Pap clan CL0125, as well as an astrovirus capsid protein profile PF03115) (101) was compared to proteins encoded by the predicted ORFs using HMMER 3.4 ‘hmmsearch’ with the ‘--max -E 0.001’ parameters. Only hits characterized by independent E-value <0.001 and covering >100 aa of the target protein sequence were considered. R packages Rhmmer 0.2.0 and IRanges 2.32.0 (102) were used to identify overlapping hits. To define clusters of hits, all hits associated with a target protein sequence were considered as graph vertices, and if two hits overlapped by >70 aa of the target protein sequence we added an edge between the corresponding vertices. Connected components of such graph, found with the help of R package Igraph 2.0.3, became clusters of hits. A hit characterized by the highest bit-score within a cluster was used for annotation.

When two annotated Pro-Chy domains overlapped by >30 aa (40 cases), only the annotation characterized by the higher bit-score was preserved. To search for potentially missed Pro-Pap of *Hypofuvirales*, the sequences of 16 initially annotated *Hypofuvirales* Pro-Pap were extracted and compared to *Hypofuvirales* proteins using BLASTP 2.17.0+ with the ‘-evalue 0.001’ parameter (99). Only hits covering >80% of the query were considered, allowing to annotate 25 more *Hypofuvirales* Pro-Pap. Selected proteins were additionally analyzed using HHpred (103).

### Pro-Chy catalytic residue analysis

All 2,708 chymotrypsin-like protease domains identified in this study were aligned using Muscle 5.3 ‘Super5’ algorithm (91). 14 sequences that contained a residue other than cysteine or serine in the MSA column corresponding to the catalytic residue were excluded from consideration. The MSA was used to analyze the distribution of cysteine and serine catalytic residues in chymotrypsin-like proteases of *Stelpaviricetes*.

### Structure-based protein domain annotation

Structures of polyprotein N-termini were predicted by AlphaFold 3.0.1 with default settings (104). Besides modeling the structure of the entire polyprotein N-termini, polyprotein N-termini were sliced into 200 aa long fragments using a sliding window with a 100 aa step to improve model accuracy of individual domains. Further, predicted structures of the entire N-termini were cut manually into individual globular domains. Obtained structure predictions were compared against a local PDB70 database (2021) using a local standalone version of Dali (DaliLite, version 5.1) (105) and against the BFVD database (downloaded in 2024) (106) and the Foldseek databases ‘pdb’, ‘alphafold-proteome’ and ‘alphafold-swissprot’ (https://foldseek.steineggerlab.workers.dev/; all downloaded in 2023) using Foldseek (version d2d09b588f50d5f8e2fd7a958377a33b2f725415) (107). Hits were inspected manually including structure superposition and inspection for catalytic residues to identify reliable protease hits. Protein structures were visualized with ChimeraX 1.3 and superimposed using the ‘match’ command of ChimeraX 1.3 (108). To access conservation of the domains annotated based on structure comparison, the entire polyprotein N-termini were aligned using Muscle 5.3 (91) and the alignments were visualized using ESPript 3.2 (109).

### Analysis of distant homologs

Protein sequences of Pro-Pap, S1H, S2H and CPf domains were extracted from the viruses representing species-level vOTUs and filtered for completeness. The non-permuted papain-like protease sequences were restricted to the protease catalytic core, while capsid protein sequences were restricted to a conserved C-terminal portion. The sequences of each domain were then clustered using MMseqs2 3.0 with the ‘--cov-mode 1 -c 0.8 --min-seq-id 0.3’ parameters (93).

A representative sequence from each cluster was compared to the NCBI Protein Reference Sequences database (25 June 2026) using BLASTP 2.17.0+ with ‘-evalue 0.001-max_target_seqs 100000’ parameters (99). Representative sequences of Pro-Pap and CPf domains were also compared to the same database using PSI-BLAST 2.17.0+ with ‘-num_iterations 3 -evalue 0.001 -max_target_seqs 100000’ parameters. A single hit, characterized by the highest bit-score among hits covering >80% of a query sequence, was selected per target sequence. The associated taxonomic information was extracted using the NCBI Entrez 25.9.

Structure of a representative sequence from each Pro-Pap and CPf cluster was predicted using AlphaFold 3.0.1 (104) and compared to the PDB25 database using DaliLite 5.1 (105). Only the hits characterized by Z-score >5 were considered. Models were visualized using PyMOL 2.6.0a0.

BLASTP hits to S1H and S2H domains were clustered using MMseqs2 3.0 with the ‘--cov-mode 1 -c 0.8 --min-seq-id 0.9’ parameters (93) and one representative per cluster was retained for phylogeny reconstruction. Sequences of S1H and S2H domains were aligned using the Muscle 5.3 ‘Super5’ algorithm (91) and filtered for completeness of I and VI motifs. MSA columns containing >50% gaps were excluded from consideration. The MSAs served for phylogeny reconstruction using FastTree 2.2.0 with the ‘-lg’ parameter (95).

**Material S1.** Domain organization of the *Stelpaviricetes* contigs. One genome per family-level vOTU is presented. For details regarding the contig maps, see legend of Fig. 2. PLA2, phospholipase A2 homolog. Shimodaira-Hasegawa (SH) test support values are indicated by white, grey and black circles on the RdRP-based phylogenetic tree.

**Material S2.** Domain organization of the contigs belonging to family *Astroviridae*. One genome per species-level vOTU is presented. For details regarding the contig maps, see legend of Fig. 2. Names of the contigs representing vOTUs that include viruses classified by ICTV are colored according to ICTV genus (see Fig. 4). Corresponding clades are indicated by branch color up to the most recent common ancestor of these vOTUs.

**Material S3.** Domain organization of the contigs belonging to families *Potyviridae* and *Potyliviridae*. The *Potyliviridae* clade is highlighted in grey. One genome per species-level vOTU is presented. For details regarding the genome maps, see legend of Fig. 2. Names of the contigs representing vOTUs that include viruses classified by ICTV are colored according to ICTV genus (see Fig. 5). Corresponding clades are indicated by branch color up to the most recent common ancestor of these vOTUs. Sequence of the second genome segment is unavailable for bymovirus LC038189.1 representing a vOTU, LC534638.1 and LC534640.1 are presented instead. Sequence of the second genome segment is unavailable for bymovirus SG001_421 representing a vOTU, MH428827.1 and MH428828.1 are presented instead.

**Material S4.** Domain organization of the contigs belonging to families *Hypo-, Parahypo- and Fusariviridae*. The *Parahypoviridae* and *Fusariviridae* clades are highlighted in grey, while the *Hypoviridae* clade is shown on white background. One genome per species-level vOTU is presented. For details regarding the genome maps, see legend of Fig. 2. Names of the contigs representing vOTUs that include viruses classified by ICTV are colored according to ICTV genus (see Fig. 6). Corresponding clades are indicated by branch color up to the most recent common ancestor of these vOTUs. Note that the genus *Iotahypovirus* is non-monophyletic.

**Material S5.** Domain organization of viruses outside of the recognized *Stelpaviricetes* families. Each page corresponds to a family-level vOTU. Family-level vOTUs composed of a single species-level vOTU are excluded; family-level vOTUs corresponding to six families recognized by ICTV are excluded. One representative per species-level vOTU is presented. For details regarding the genome maps, see legend of Fig. 2.

**Table S1.** HMMER-based taxonomic assignments made for coding-complete RNA virus genomes listed in the ICTV MSL 41.1.

**Table S2.** Information about the *Stelpaviricetes* contigs analyzed in this study.

**Table S3.** Tree topology testing results.

**Table S4.** EVEs linked to *Stelpaviricetes* contigs.

**Table S5.** Putative papain-like protease domains of celaviruses and potyliviruses.

---

**Fig. S1.**
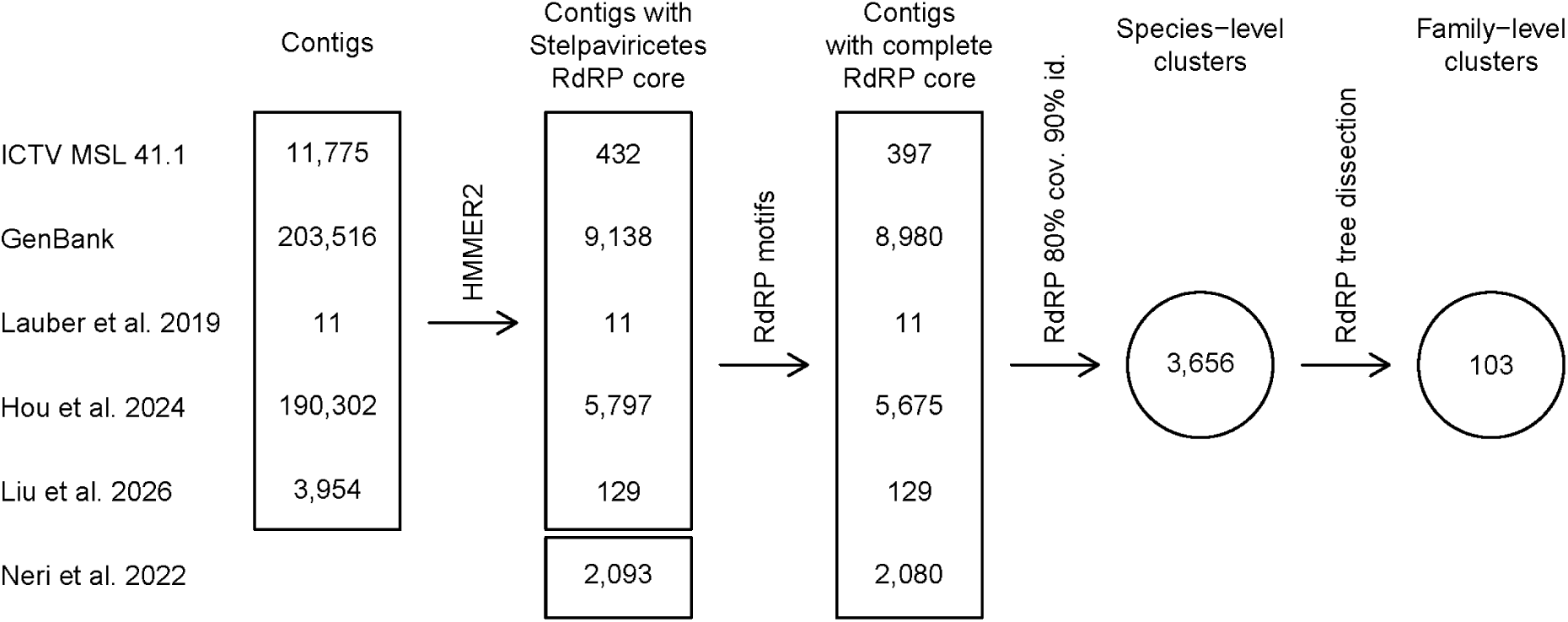
*Stelpaviricetes* contigs detection and clustering workflow.

**Fig. S2.**
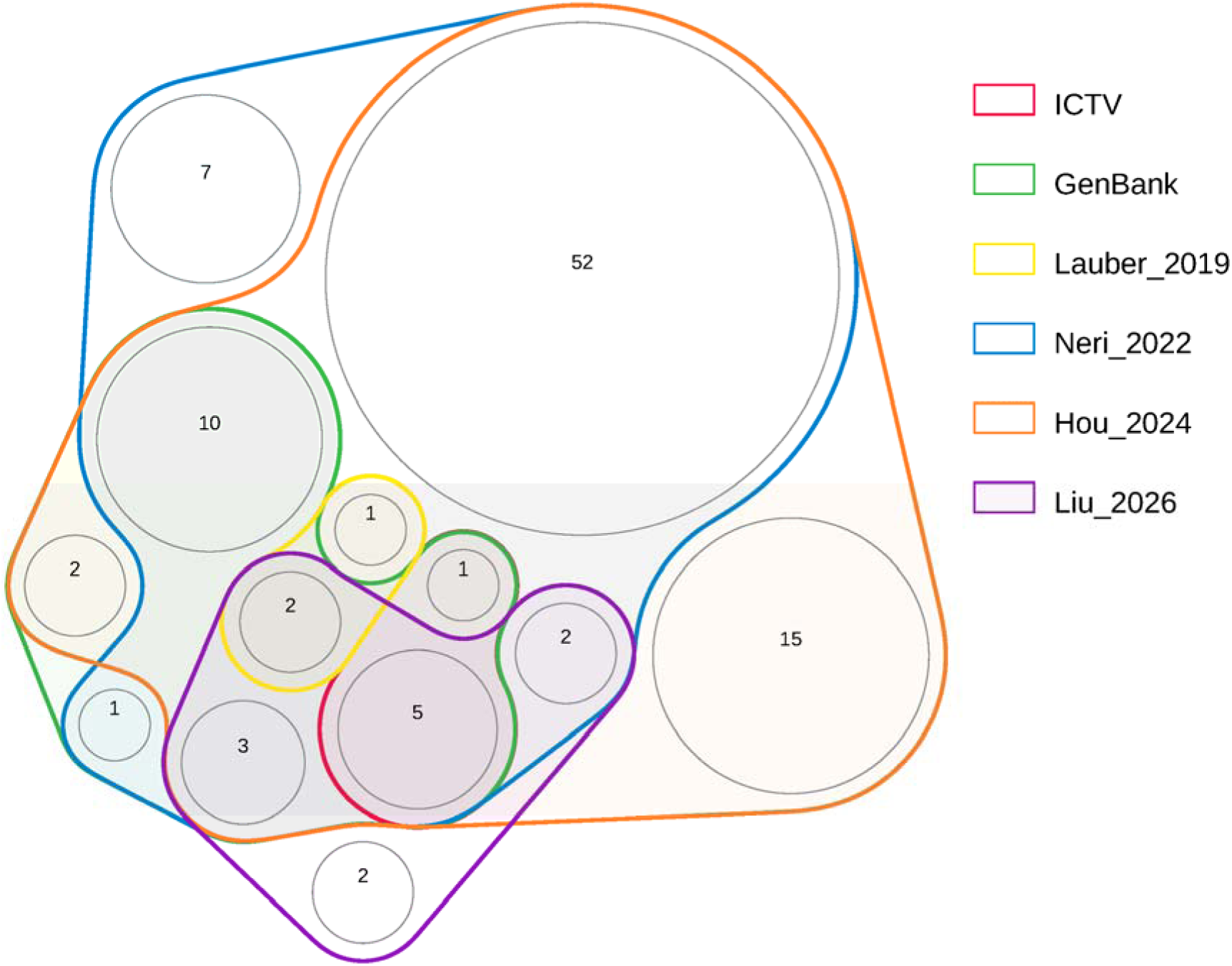
Venn diagram quantifying family-level vOTUs containing contigs from different datasets. Prepared using R package nVennR 0.2.3.

**Fig. S3.**
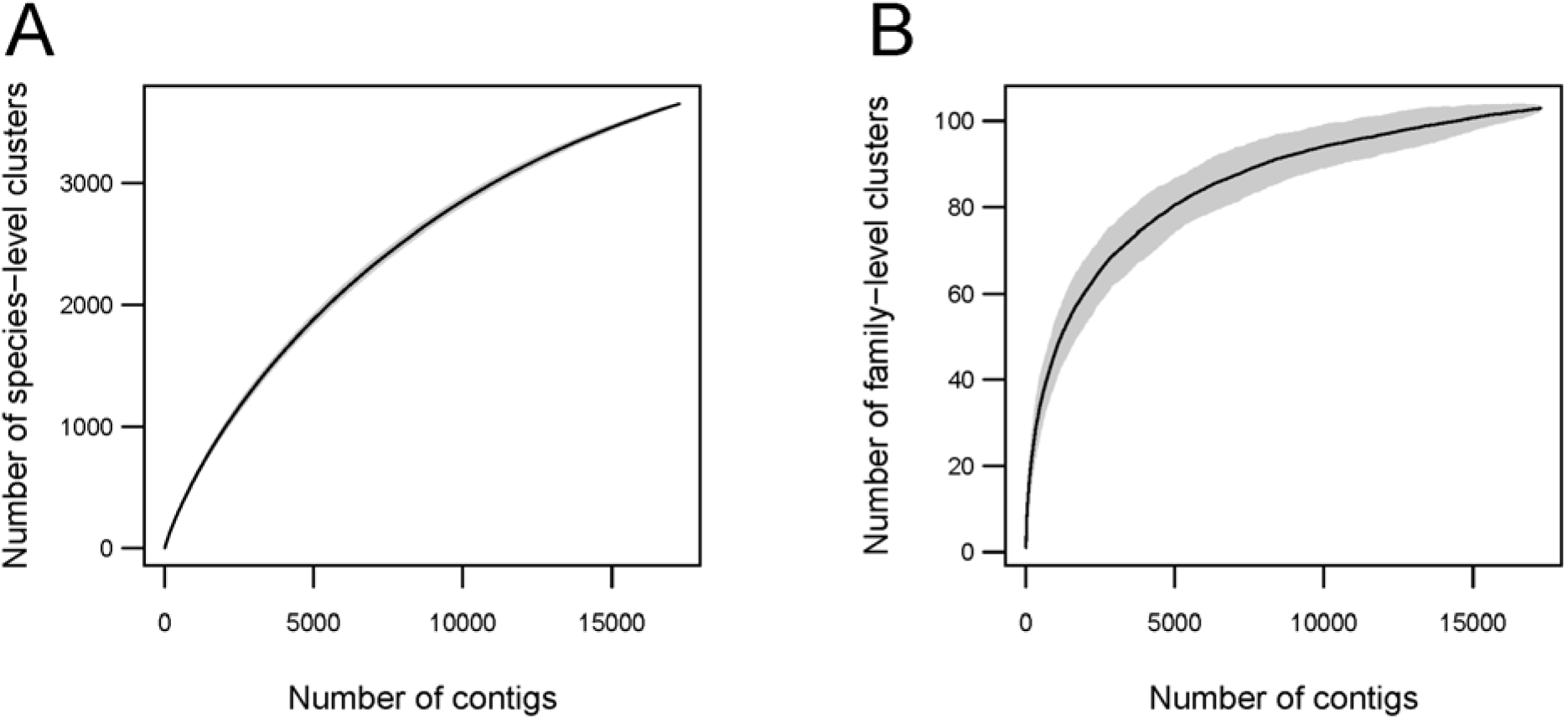
Accumulation curves for species- and family-level vOTUs. Confidence intervals are shown in grey.

**Figure S4.**
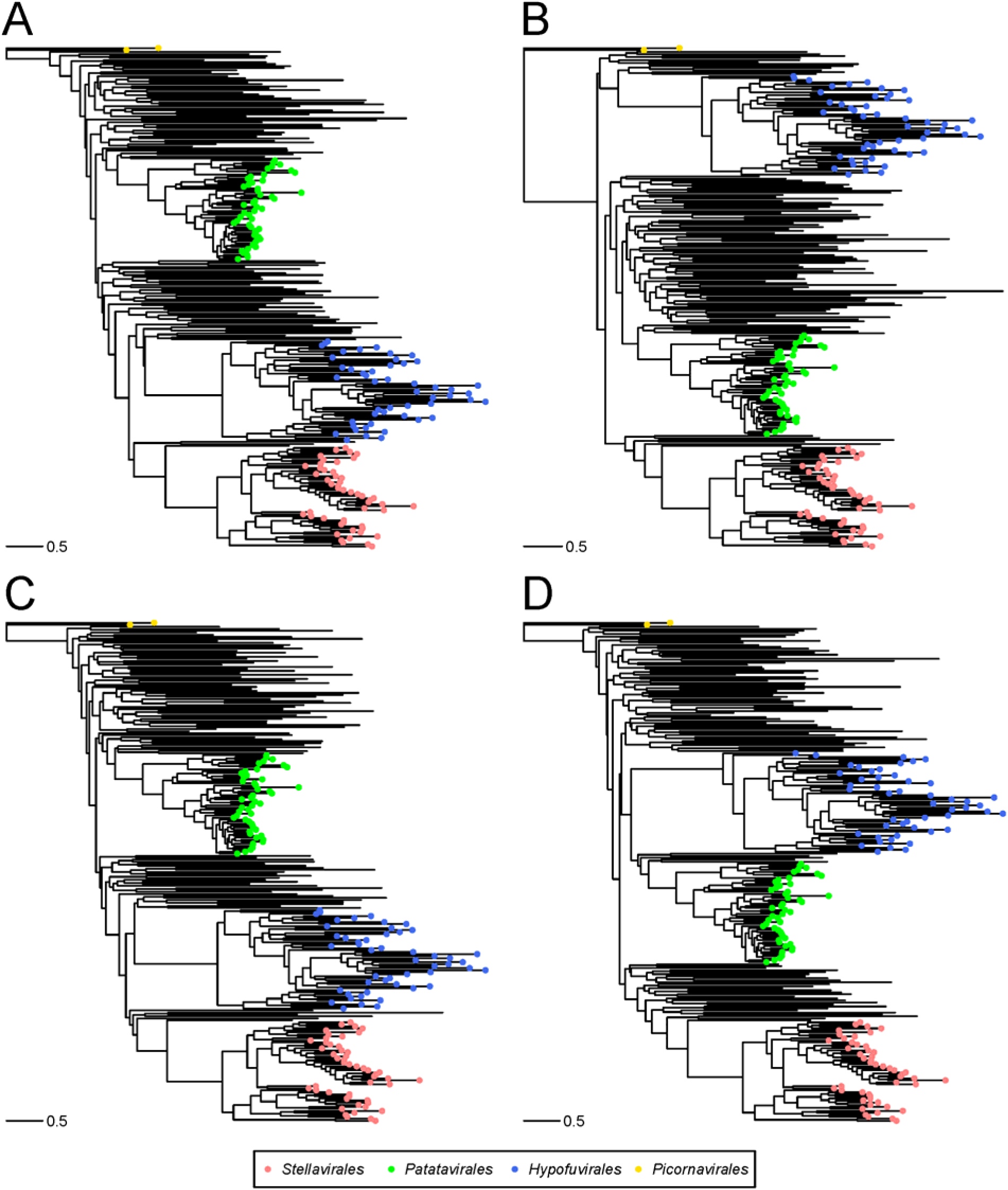
RdRP-based phylogenetic trees utilized in tree topology testing. (A) A tree reconstructed without imposing a constraint on its topology. Trees constrained to impose evolutionary affinity of (B) orders *Stellavirales* and *Patatavirales*, (C) orders *Stellavirales* and *Hypofuvirales*, (D) *Patatavirales* and *Hypofuvirales*.

**Fig. S5.**
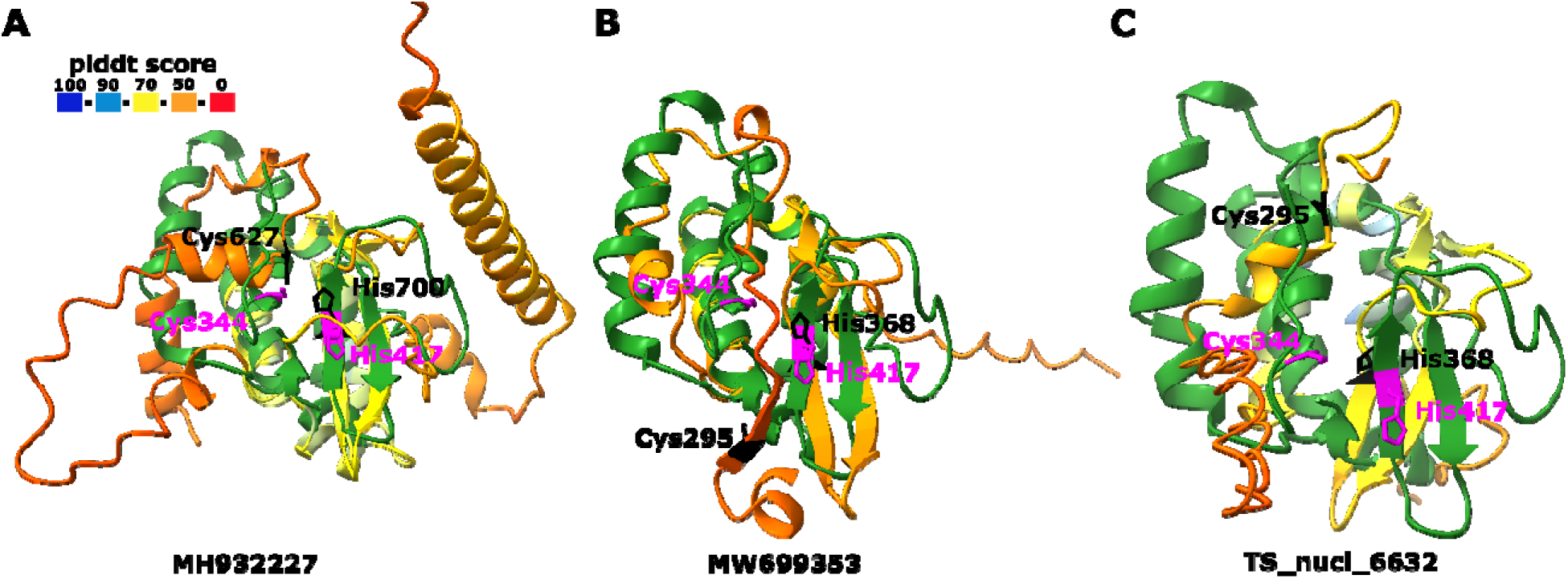
Superposition of the putative papain-like protease domain of three celaviruses with the turnip mosaic virus papain-like protease domain (PDB 3RNV). Celavirus protease models are colored according to the predicted local distance difference test (pLDDT) score, a measure of AlphaFold confidence, with catalytic dyad shown in black. The 3RNV structure is shown in green with catalytic dyad (Cys344 and His417) in magenta. (A) MH932227: aa 601-800, (B) MW699353: aa 290-426 and (C) TS_nucl_6632: aa 287-400.

**Fig. S6.**
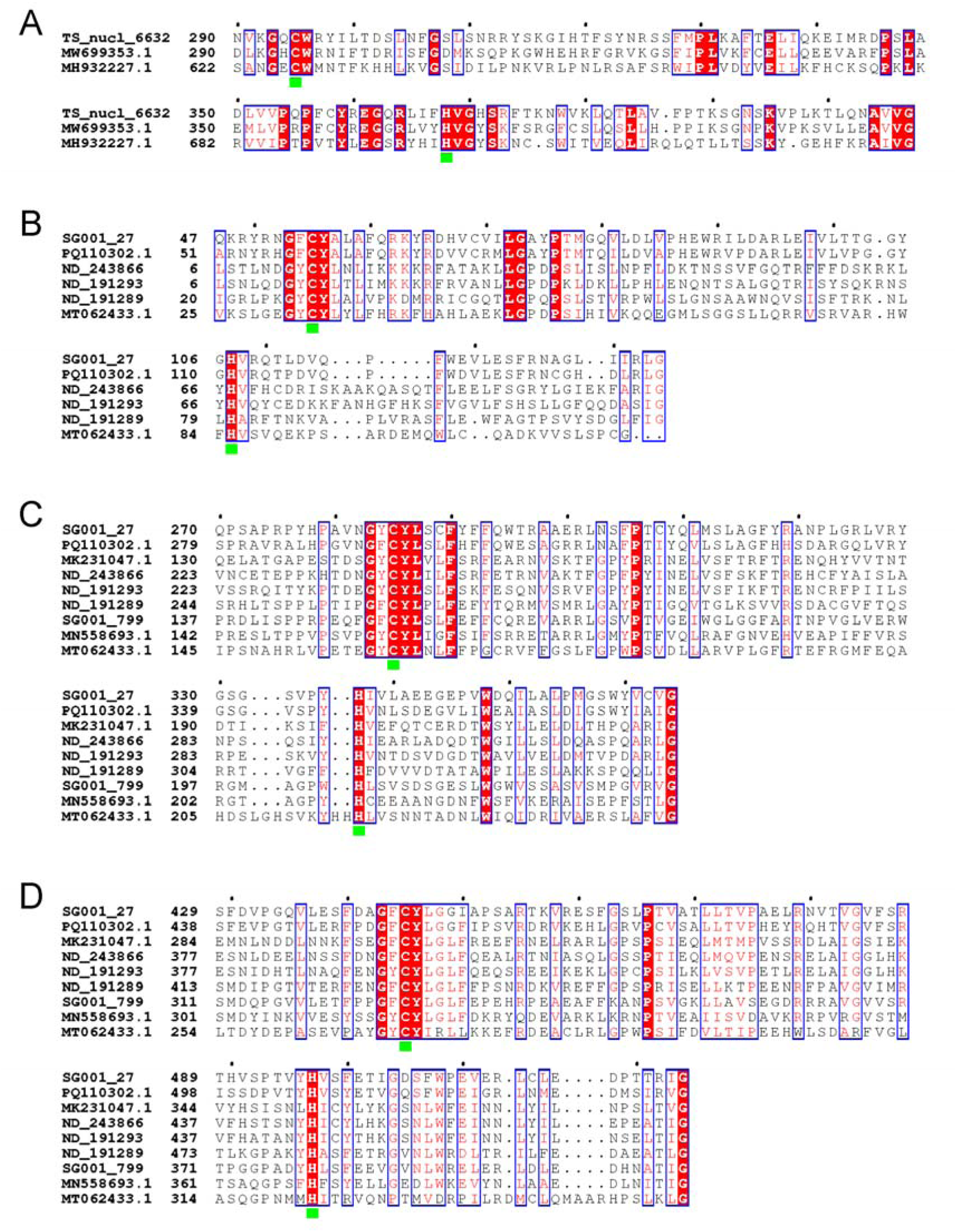
Polyprotein N-termini alignments fragments. (A) Putative protease domain of celaviruses. (B) First, (C) second and (D) third putative protease domains of the potyliviruses. Absolutely conserved residues are shown on red background with a blue frame, partially conserved residues – in red font with a blue frame. Putative catalytic residues are indicated by green squares. At the beginning of each block of sequences, a polyprotein residue number is specified for every sequence.

**Fig. S7.**
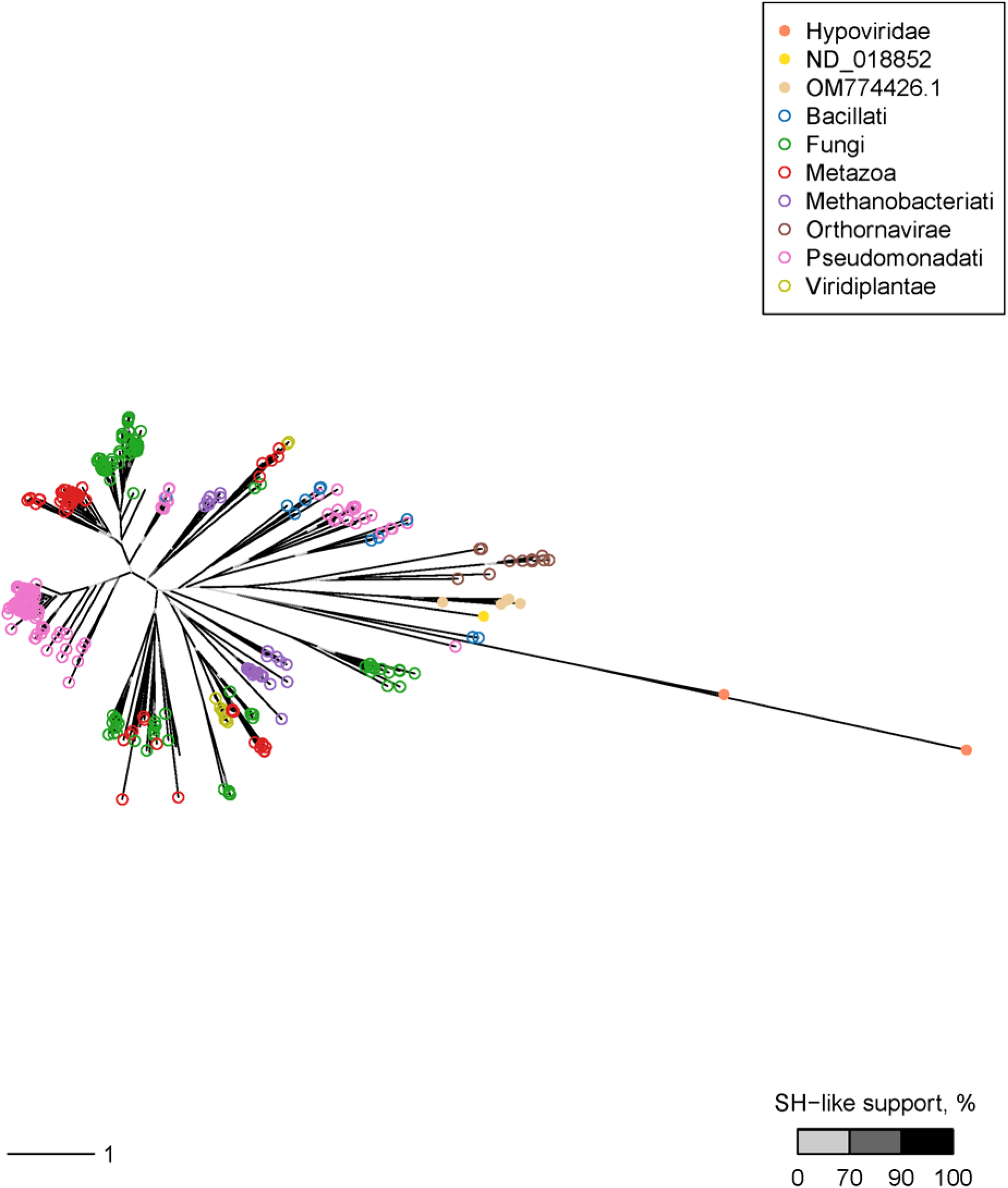
S1H-based phylogenetic tree. *Stelpaviricetes* tips are indicated by filled dots colored according to family-level vOTU, non-*Stelpaviricetes* tips are indicated by empty dots colored according to kingdom. Color of the internal branches indicates SH-like support values; terminal branches are colored in black.

**Fig. S8.**
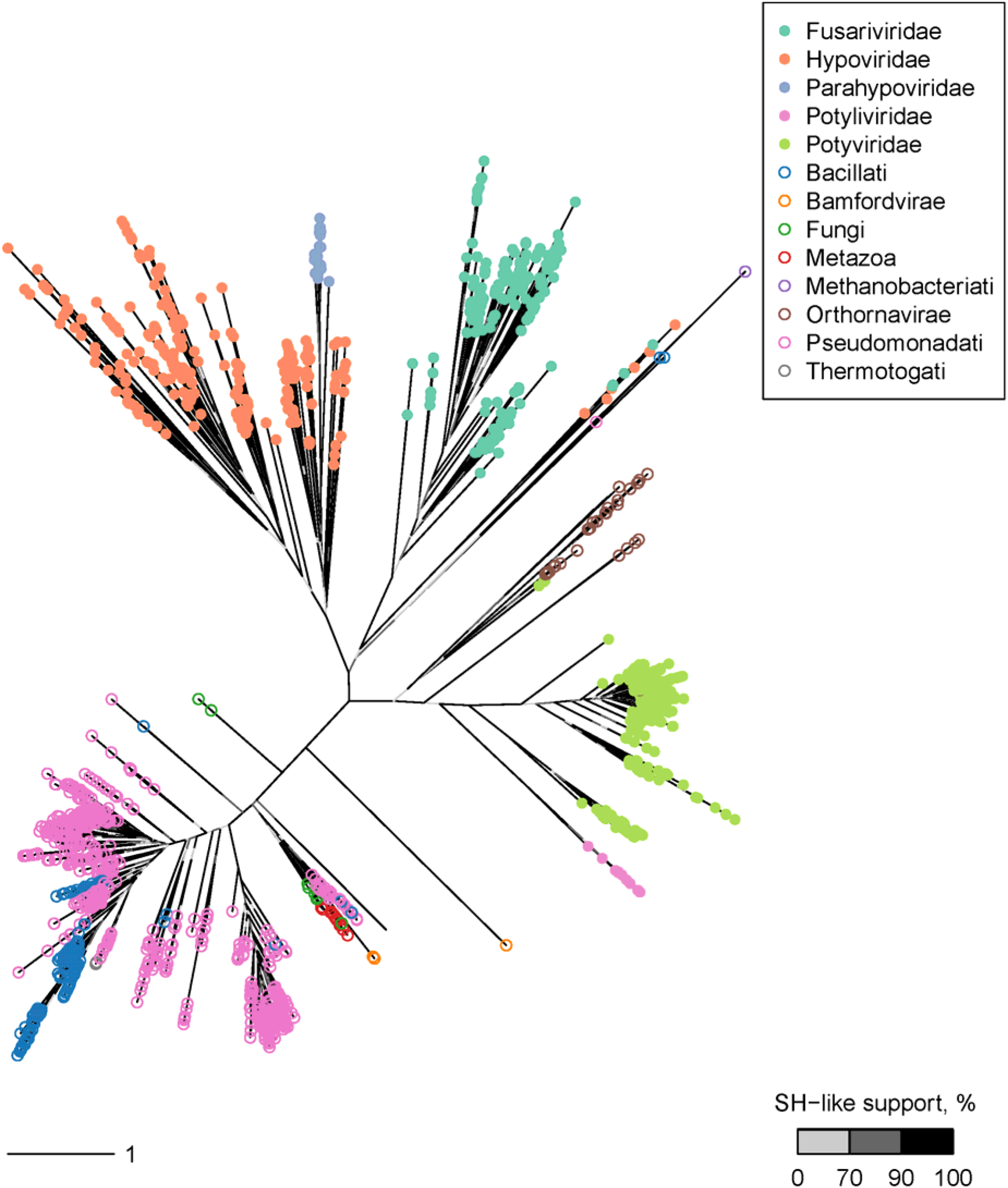
S2H-based phylogenetic tree. *Stelpaviricetes* tips are indicated by filled dots colored according to family-level vOTU, non-*Stelpaviricetes* tips are indicated by empty dots colored according to kingdom. Color of the internal branches indicates SH-like support values; terminal branches are colored in black.

## REFERENCES

1. Paez-Espino D, Eloe-Fadrosh EA, Pavlopoulos GA, Thomas AD, Huntemann M, Mikhailova N, Rubin E, Ivanova NN, Kyrpides NC. 2016. Uncovering Earth’s virome. Nature 536:425–30.

2. Gregory AC, Zayed AA, Conceicao-Neto N, Temperton B, Bolduc B, Alberti A, Ardyna M, Arkhipova K, Carmichael M, Cruaud C, Dimier C, Dominguez-Huerta G, Ferland J, Kandels S, Liu Y, Marec C, Pesant S, Picheral M, Pisarev S, Poulain J, Tremblay JE, Vik D, Tara Oceans C, Babin M, Bowler C, Culley AI, de Vargas C, Dutilh BE, Iudicone D, Karp-Boss L, Roux S, Sunagawa S, Wincker P, Sullivan MB. 2019. Marine DNA Viral Macro- and Microdiversity from Pole to Pole. Cell 177:1109–1123 e14.

3. Neri U, Wolf YI, Roux S, Camargo AP, Lee B, Kazlauskas D, Chen IM, Ivanova N, Zeigler Allen L, Paez-Espino D, Bryant DA, Bhaya D, Consortium RNAVD, Krupovic M, Dolja VV, Kyrpides NC, Koonin EV, Gophna U. 2022. Expansion of the global RNA virome reveals diverse clades of bacteriophages. Cell 185:4023–4037 e18.

4. Edgar RC, Taylor B, Lin V, Altman T, Barbera P, Meleshko D, Lohr D, Novakovsky G, Buchfink B, Al-Shayeb B, Banfield JF, de la Pena M, Korobeynikov A, Chikhi R, Babaian A. 2022. Petabase-scale sequence alignment catalyses viral discovery. Nature 602:142–147.

5. Hou X, He Y, Fang P, Mei SQ, Xu Z, Wu WC, Tian JH, Zhang S, Zeng ZY, Gou QY, Xin GY, Le SJ, Xia YY, Zhou YL, Hui FM, Pan YF, Eden JS, Yang ZH, Han C, Shu YL, Guo D, Li J, Holmes EC, Li ZR, Shi M. 2024. Using artificial intelligence to document the hidden RNA virosphere. Cell 187:6929–6942 e16.

6. Dolja VV, Koonin EV. 2018. Metagenomics reshapes the concepts of RNA virus evolution by revealing extensive horizontal virus transfer. Virus Res 244:36–52.

7. Zhang YZ, Chen YM, Wang W, Qin XC, Holmes EC. 2019. Expanding the RNA Virosphere by Unbiased Metagenomics. Annu Rev Virol 6:119–139.

8. Dion MB, Oechslin F, Moineau S. 2020. Phage diversity, genomics and phylogeny. Nat Rev Microbiol 18:125–138.

9. Krupovic M, Dolja VV, Koonin EV. 2019. Origin of viruses: primordial replicators recruiting capsids from hosts. Nat Rev Microbiol 17:449–458.

10. Krupovic M, Dolja VV, Koonin EV. 2020. The LUCA and its complex virome. Nat Rev Microbiol 18:661–670.

11. Koonin EV, Dolja VV, Krupovic M. 2022. The logic of virus evolution. Cell Host Microbe 30:917–929.

12. Krupovic M, Dolja VV, Koonin EV. 2023. The virome of the last eukaryotic common ancestor and eukaryogenesis. Nat Microbiol 8:1008–1017.

13. Koonin EV, Dolja VV, Krupovic M, Varsani A, Wolf YI, Yutin N, Zerbini FM, Kuhn JH. 2020. Global Organization and Proposed Megataxonomy of the Virus World. Microbiol Mol Biol Rev 84.

14. International Committee on Taxonomy of Viruses Executive C. 2020. The new scope of virus taxonomy: partitioning the virosphere into 15 hierarchical ranks. Nat Microbiol 5:668–674.

15. Wolf YI, Kazlauskas D, Iranzo J, Lucia-Sanz A, Kuhn JH, Krupovic M, Dolja VV, Koonin EV. 2018. Origins and Evolution of the Global RNA Virome. mBio 9.

16. Krupovic M, Blomberg J, Coffin JM, Dasgupta I, Fan H, Geering AD, Gifford R, Harrach B, Hull R, Johnson W, Kreuze JF, Lindemann D, Llorens C, Lockhart B, Mayer J, Muller E, Olszewski NE, Pappu HR, Pooggin MM, Richert-Poggeler KR, Sabanadzovic S, Sanfacon H, Schoelz JE, Seal S, Stavolone L, Stoye JP, Teycheney PY, Tristem M, Koonin EV, Kuhn JH. 2018. Ortervirales: New Virus Order Unifying Five Families of Reverse-Transcribing Viruses. J Virol 92.

17. Koonin EV, Kuhn JH, Dolja VV, Krupovic M. 2024. Megataxonomy and global ecology of the virosphere. ISME J 18.

18. Urayama SI, Fukudome A, Hirai M, Okumura T, Nishimura Y, Takaki Y, Kurosawa N, Koonin EV, Krupovic M, Nunoura T. 2024. Double-stranded RNA sequencing reveals distinct riboviruses associated with thermoacidophilic bacteria from hot springs in Japan. Nat Microbiol 9:514–523.

19. Neuman BW, Smart A, Gilmer O, Smyth RP, Vaas J, Boker N, Samborskiy DV, Bartenschlager R, Seitz S, Gorbalenya AE, Caliskan N, Lauber C. 2025. Giant RNA genomes: Roles of host, translation elongation, genome architecture, and proteome in nidoviruses. Proc Natl Acad Sci U S A 122:e2413675122.

20. Koonin EV, Dolja VV, Krupovic M, Kuhn JH. 2021. Viruses Defined by the Position of the Virosphere within the Replicator Space. Microbiol Mol Biol Rev 85:e0019320.

21. Forgia M, Navarro B, Daghino S, Cervera A, Gisel A, Perotto S, Aghayeva DN, Akinyuwa MF, Gobbi E, Zheludev IN, Edgar RC, Chikhi R, Turina M, Babaian A, Di Serio F, de la Pena M. 2023. Hybrids of RNA viruses and viroid-like elements replicate in fungi. Nat Commun 14:2591.

22. Lee BD, Neri U, Roux S, Wolf YI, Camargo AP, Krupovic M, Consortium RNAVD, Simmonds P, Kyrpides N, Gophna U, Dolja VV, Koonin EV. 2023. Mining metatranscriptomes reveals a vast world of viroid-like circular RNAs. Cell 186:646–661 e4.

23. Cortez V, Meliopoulos VA, Karlsson EA, Hargest V, Johnson C, Schultz-Cherry S. 2017. Astrovirus Biology and Pathogenesis. Annu Rev Virol 4:327–348.

24. Wohlgemuth N, Honce R, Schultz-Cherry S. 2019. Astrovirus evolution and emergence. Infect Genet Evol 69:30–37.

25. Yang X, Li Y, Wang A. 2021. Research Advances in Potyviruses: From the Laboratory Bench to the Field. Annu Rev Phytopathol 59:1–29.

26. Pollari ME, Aspelin WWE, Wang L, Makinen KM. 2024. The Molecular Maze of Potyviral and Host Protein Interactions. Annu Rev Virol 11:147–170.

27. Jiang B, Monroe SS, Koonin EV, Stine SE, Glass RI. 1993. RNA sequence of astrovirus: distinctive genomic organization and a putative retrovirus-like ribosomal frameshifting signal that directs the viral replicase synthesis. Proc Natl Acad Sci U S A 90:10539–43.

28. York RL, Yousefi PA, Bogdanoff W, Haile S, Tripathi S, DuBois RM. 2015. Structural, Mechanistic, and Antigenic Characterization of the Human Astrovirus Capsid. J Virol 90:2254–63.

29. Dolja VV, Boyko VP, Agranovsky AA, Koonin EV. 1991. Phylogeny of capsid proteins of rod-shaped and filamentous RNA plant viruses: two families with distinct patterns of sequence and probably structure conservation. Virology 184:79–86.

30. Agirrezabala X, Mendez-Lopez E, Lasso G, Sanchez-Pina MA, Aranda M, Valle M. 2015. The near-atomic cryoEM structure of a flexible filamentous plant virus shows homology of its coat protein with nucleoproteins of animal viruses. Elife 4:e11795.

31. Lefkowitz E, Hendrickson R. 2026. ICTV Master Species List, 41 ed doi:10.5281/zenodo.19154110.

32. Koonin EV, Choi GH, Nuss DL, Shapira R, Carrington JC. 1991. Evidence for common ancestry of a chestnut blight hypovirulence-associated double-stranded RNA and a group of positive-strand RNA plant viruses. Proc Natl Acad Sci U S A 88:10647–51.

33. Simmonds P, Adriaenssens EM, Lefkowitz EJ, Oksanen HM, Zerbini FM, Alfenas-Zerbini P, Aylward FO, Dempsey DM, Freitas-Astua J, Hendrickson RC, Hughes HR, Krupovic M, Kuhn JH, Lobocka M, Mayne R, Mushegian AR, Penzes JJ, Reyes Munoz A, Robertson DL, Roux S, Rubino L, Sabanadzovic S, Smith DB, Suzuki N, Turner D, Doorslaer KV, Varsani A. 2025. Changes to virus taxonomy, the international code of virus classification and nomenclature, and the ICTV statutes ratified by the International Committee on Taxonomy of Viruses (2025). Arch Virol 171:23.

34. Sayers EW, Cavanaugh M, Frisse L, Pruitt KD, Schneider VA, Underwood BA, Yankie L, Karsch-Mizrachi I. 2025. GenBank 2025 update. Nucleic Acids Res 53:D56–D61.

35. Lauber C, Seifert M, Bartenschlager R, Seitz S. 2019. Discovery of highly divergent lineages of plant-associated astro-like viruses sheds light on the emergence of potyviruses. Virus Res 260:38–48.

36. Liu Q, Gong Z, Koonin EV, Dolja VV, Han GZ. 2026. The Origins and Evolution of the RNA Virome in Land Flora. Mol Plant doi:10.1016/j.molp.2026.07.023.

37. Shapira R, Choi GH, Nuss DL. 1991. Virus-like genetic organization and expression strategy for a double-stranded RNA genetic element associated with biological control of chestnut blight. EMBO J 10:731–9.

38. Smart CD, Yuan W, Foglia R, Nuss DL, Fulbright DW, Hillman BI. 1999. Cryphonectria hypovirus 3, a virus species in the family hypoviridae with a single open reading frame. Virology 265:66–73.

39. Zhang R, Liu S, Chiba S, Kondo H, Kanematsu S, Suzuki N. 2014. A novel single-stranded RNA virus isolated from a phytopathogenic filamentous fungus, Rosellinia necatrix, with similarity to hypo-like viruses. Front Microbiol 5:360.

40. Rose H, Doring I, Vetten HJ, Menzel W, Richert-Poggeler KR, Maiss E. 2019. Complete genome sequence and construction of an infectious full-length cDNA clone of celery latent virus - an unusual member of a putative new genus within the Potyviridae. J Gen Virol 100:308–320.

41. Wolf YI, Silas S, Wang Y, Wu S, Bocek M, Kazlauskas D, Krupovic M, Fire A, Dolja VV, Koonin EV. 2020. Doubling of the known set of RNA viruses by metagenomic analysis of an aquatic virome. Nat Microbiol 5:1262–1270.

42. Minh BQ, Schmidt HA, Chernomor O, Schrempf D, Woodhams MD, von Haeseler A, Lanfear R. 2020. IQ-TREE 2: New Models and Efficient Methods for Phylogenetic Inference in the Genomic Era. Mol Biol Evol 37:1530–1534.

43. Kishino H, Hasegawa M. 1989. Evaluation of the maximum likelihood estimate of the evolutionary tree topologies from DNA sequence data, and the branching order in hominoidea. J Mol Evol 29:170–9.

44. Shimodaira H, Hasegawa M. 1999. Multiple Comparisons of Log-Likelihoods with Applications to Phylogenetic Inference. Molecular Biology and Evolution 16:1114–1114.

45. Strimmer K, Rambaut A. 2002. Inferring confidence sets of possibly misspecified gene trees. Proc Biol Sci 269:137–42.

46. Shimodaira H. 2002. An approximately unbiased test of phylogenetic tree selection. Syst Biol 51:492–508.

47. Shi M, Lin XD, Tian JH, Chen LJ, Chen X, Li CX, Qin XC, Li J, Cao JP, Eden JS, Buchmann J, Wang W, Xu J, Holmes EC, Zhang YZ. 2016. Redefining the invertebrate RNA virosphere. Nature 540:539–543.

48. Huang HJ, Ye ZX, Wang X, Yan XT, Zhang Y, He YJ, Qi YH, Zhang XD, Zhuo JC, Lu G, Lu JB, Mao QZ, Sun ZT, Yan F, Chen JP, Zhang CX, Li JM. 2021. Diversity and infectivity of the RNA virome among different cryptic species of an agriculturally important insect vector: whitefly Bemisia tabaci. NPJ Biofilms Microbiomes 7:43.

49. Li LL, Ye ZX, Chen JP, Zhang CX, Huang HJ, Li JM. 2022. Characterization of Two Novel Insect-Specific Viruses Discovered in the Green Leafhopper, Cicadella viridis. Insects 13.

50. Hernandez-Pelegrin L, Rodriguez-Gomez A, Abelaira AB, Reche MC, Crava C, Lim FS, Bielza P, Herrero S. 2024. Rich diversity of RNA viruses in the biological control agent, Orius laevigatus. J Invertebr Pathol 206:108175.

51. Becerra-Garcia RE, Hernandez-Pelegrin L, Crava CM, Herrero S. 2025. Characterization of the Tuta absoluta virome reveals higher viral diversity in field populations. J Invertebr Pathol 211:108340.

52. Shi M, Lin XD, Chen X, Tian JH, Chen LJ, Li K, Wang W, Eden JS, Shen JJ, Liu L, Holmes EC, Zhang YZ. 2018. The evolutionary history of vertebrate RNA viruses. Nature 556:197–202.

53. Yang K, Shen W, Li Y, Li Z, Miao W, Wang A, Cui H. 2019. Areca Palm Necrotic Ringspot Virus, Classified Within a Recently Proposed Genus Arepavirus of the Family Potyviridae, Is Associated With Necrotic Ringspot Disease in Areca Palm. Phytopathology 109:887–894.

54. Verchot J, Koonin EV, Carrington JC. 1991. The 35-kDa protein from the N-terminus of the potyviral polyprotein functions as a third virus-encoded proteinase. Virology 185:527–35.

55. Rohozkova J, Navratil M. 2011. P1 peptidase--a mysterious protein of family Potyviridae. J Biosci 36:189–200.

56. Valli A, Martin-Hernandez AM, Lopez-Moya JJ, Garcia JA. 2006. RNA silencing suppression by a second copy of the P1 serine protease of Cucumber vein yellowing ipomovirus, a member of the family Potyviridae that lacks the cysteine protease HCPro. J Virol 80:10055–63.

57. Berman HM, Westbrook J, Feng Z, Gilliland G, Bhat TN, Weissig H, Shindyalov IN, Bourne PE. 2000. The Protein Data Bank. Nucleic Acids Res 28:235–42.

58. Chiapello M, Rodriguez-Romero J, Ayllon MA, Turina M. 2020. Analysis of the virome associated to grapevine downy mildew lesions reveals new mycovirus lineages. Virus Evol 6:veaa058.

59. Chiapello M, Rodríguez-Romero J, Nerva L, Forgia M, Chitarra W, Ayllón MA, Turina M. 2020. Putative new plant viruses associated with Plasmopara viticola-infected grapevine samples. Annals of Applied Biology 176:180–191.

60. Wang J, Ni Y, Liu X, Zhao H, Xiao Y, Xiao X, Li S, Liu H. 2021. Divergent RNA viruses in Macrophomina phaseolina exhibit potential as virocontrol agents. Virus Evol 7:veaa095.

61. Jo Y, Choi H, Chu H, Cho WK. 2022. Unveiling Mycoviromes Using Fungal Transcriptomes. Int J Mol Sci 23.

62. Abdoulaye AH, Hai D, Tang Q, Jiang D, Fu Y, Cheng J, Lin Y, Li B, Kotta-Loizou I, Xie J. 2021. Two distant helicases in one mycovirus: evidence of horizontal gene transfer between mycoviruses, coronaviruses and other nidoviruses. Virus Evol 7:veab043.

63. Murolo S, De Miccolis Angelini RM, Faretra F, Romanazzi G. 2018. Phenotypic and Molecular Investigations on Hypovirulent Cryphonectria parasitica in Italy. Plant Dis 102:540–545.

64. Iyer LM, Koonin EV, Aravind L. 2004. Novel predicted peptidases with a potential role in the ubiquitin signaling pathway. Cell Cycle 3:1440–50.

65. Abdoulaye AH, Jia J, Abbas A, Hai D, Cheng J, Fu Y, Lin Y, Jiang D, Xie J. 2022. Fusarivirus accessory helicases present an evolutionary link for viruses infecting plants and fungi. Virol Sin 37:427–436.

66. Chiba S, Suzuki N, Velasco L, Ayllon MA, Lee-Marzano SY, Sun L, Sabanadzovic S, Turina M. 2024. ICTV Virus Taxonomy Profile: Fusariviridae 2024. J Gen Virol 105.

67. Wang M, Wang Y, Sun X, Cheng J, Fu Y, Liu H, Jiang D, Ghabrial SA, Xie J. 2015. Characterization of a Novel Megabirnavirus from Sclerotinia sclerotiorum Reveals Horizontal Gene Transfer from Single-Stranded RNA Virus to Double-Stranded RNA Virus. J Virol 89:8567–79.

68. Xue F, Sun Y, Yan L, Zhao C, Chen J, Bartlam M, Li X, Lou Z, Rao Z. 2010. The crystal structure of porcine reproductive and respiratory syndrome virus nonstructural protein Nsp1beta reveals a novel metal-dependent nuclease. J Virol 84:6461–71.

69. Guo B, Lin J, Ye K. 2011. Structure of the autocatalytic cysteine protease domain of potyvirus helper-component proteinase. J Biol Chem 286:21937–43.

70. Suh HY, Kim JH, Woo JS, Ku B, Shin EJ, Yun Y, Oh BH. 2012. Crystal structure of DeSI-1, a novel deSUMOylase belonging to a putative isopeptidase superfamily. Proteins 80:2099–104.

71. Hersch SJ, Watanabe N, Stietz MS, Manera K, Kamal F, Burkinshaw B, Lam L, Pun A, Li M, Savchenko A, Dong TG. 2020. Envelope stress responses defend against type six secretion system attacks independently of immunity proteins. Nat Microbiol 5:706–714.

72. Luebben AV, Bender D, Becker S, Crowther LM, Erven I, Hofmann K, Soding J, Klemp H, Bellotti C, Stauble A, Qiu T, Kathayat RS, Dickinson BC, Gartner J, Sheldrick GM, Kratzner R, Steinfeld R. 2022. Cln5 represents a new type of cysteine-based S-depalmitoylase linked to neurodegeneration. Sci Adv 8:eabj8633.

73. Chase O, Javed A, Byrne MJ, Thuenemann EC, Lomonossoff GP, Ranson NA, Lopez-Moya JJ. 2023. CryoEM and stability analysis of virus-like particles of potyvirus and ipomovirus infecting a common host. Commun Biol 6:433.

74. DiMaio F, Chen CC, Yu X, Frenz B, Hsu YH, Lin NS, Egelman EH. 2015. The molecular basis for flexibility in the flexible filamentous plant viruses. Nat Struct Mol Biol 22:642–4.

75. Olal D, Dick A, Woods VL, Jr., Liu T, Li S, Devignot S, Weber F, Saphire EO, Daumke O. 2014. Structural insights into RNA encapsidation and helical assembly of the Toscana virus nucleoprotein. Nucleic Acids Res 42:6025–37.

76. Zamora M, Mendez-Lopez E, Agirrezabala X, Cuesta R, Lavin JL, Sanchez-Pina MA, Aranda MA, Valle M. 2017. Potyvirus virion structure shows conserved protein fold and RNA binding site in ssRNA viruses. Sci Adv 3:eaao2182.

77. Sabanadzovic S, Aboughanem-Sabanadzovic N, Turina M, Suzuki N, Krupovic M. 2026. "Tobaliviridae", a new family of filamentous mycoviruses in the order Martellivirales. Arch Virol 171:50.

78. Mutz P, Camargo AP, Sahakyan H, Neri U, Butkovic A, Wolf YI, Krupovic M, Dolja VV, Koonin EV. 2025. The protein structurome of Orthornavirae and its dark matter. mBio 16:e0320024.

79. Gorbalenya AE, Koonin EV. 1989. Viral proteins containing the purine NTP-binding sequence pattern. Nucleic Acids Res 17:8413–40.

80. Klawonn I, Van den Wyngaert S, Parada AE, Arandia-Gorostidi N, Whitehouse MJ, Grossart HP, Dekas AE. 2021. Characterizing the "fungal shunt": Parasitic fungi on diatoms affect carbon flow and bacterial communities in aquatic microbial food webs. Proc Natl Acad Sci U S A 118.

81. Genre A, Lanfranco L, Perotto S, Bonfante P. 2020. Unique and common traits in mycorrhizal symbioses. Nat Rev Microbiol 18:649–660.

82. Hong S, Shang J, Sun Y, Tang G, Wang C. 2024. Fungal infection of insects: molecular insights and prospects. Trends Microbiol 32:302–316.

83. Szantho LL, Merenyi Z, Donoghue P, Gabaldon T, Nagy LG, Szollosi GJ, Ocana-Pallares E. 2025. A timetree of Fungi dated with fossils and horizontal gene transfers. Nat Ecol Evol 9:1989–2001.

84. Xie J, Jiang D. 2024. Understanding the Diversity, Evolution, Ecology, and Applications of Mycoviruses. Annu Rev Microbiol 78:595–620.

85. Dolja VV, Krupovic M, Koonin EV. 2020. Deep Roots and Splendid Boughs of the Global Plant Virome. Annu Rev Phytopathol 58:23–53.

86. Rodamilans B, Rincón Barrado M, Cobos Piñuela A, Simón Mateo C, Valli AA. 2026. Discovery of novel members of the Potyviridae family reveals expanded diversity, a broad host range, and evidence of fungal and oomycete infections. bioRxiv doi:10.64898/2026.05.29.728796.

87. O’Leary NA, Cox E, Holmes JB, Anderson WR, Falk R, Hem V, Tsuchiya MTN, Schuler GD, Zhang X, Torcivia J, Ketter A, Breen L, Cothran J, Bajwa H, Tinne J, Meric PA, Hlavina W, Schneider VA. 2024. Exploring and retrieving sequence and metadata for species across the tree of life with NCBI Datasets. Sci Data 11:732.

88. Grant BJ, Rodrigues AP, ElSawy KM, McCammon JA, Caves LS. 2006. Bio3d: an R package for the comparative analysis of protein structures. Bioinformatics 22:2695–6.

89. Shen W, Sipos B, Zhao L. 2024. SeqKit2: A Swiss army knife for sequence and alignment processing. Imeta 3:e191.

90. Rice P, Longden I, Bleasby A. 2000. EMBOSS: the European Molecular Biology Open Software Suite. Trends Genet 16:276–7.

91. Edgar RC. 2022. Muscle5: High-accuracy alignment ensembles enable unbiased assessments of sequence homology and phylogeny. Nat Commun 13:6968.

92. Charon J, Buchmann JP, Sadiq S, Holmes EC. 2022. RdRp-scan: A bioinformatic resource to identify and annotate divergent RNA viruses in metagenomic sequence data. Virus Evol 8:veac082.

93. Steinegger M, Soding J. 2017. MMseqs2 enables sensitive protein sequence searching for the analysis of massive data sets. Nat Biotechnol 35:1026–1028.

94. Waterhouse AM, Procter JB, Martin DM, Clamp M, Barton GJ. 2009. Jalview Version 2--a multiple sequence alignment editor and analysis workbench. Bioinformatics 25:1189–91.

95. Price MN, Dehal PS, Arkin AP. 2010. FastTree 2--approximately maximum-likelihood trees for large alignments. PLoS One 5:e9490.

96. Paradis E, Schliep K. 2019. ape 5.0: an environment for modern phylogenetics and evolutionary analyses in R. Bioinformatics 35:526–528.

97. Revell LJ. 2024. phytools 2.0: an updated R ecosystem for phylogenetic comparative methods (and other things). PeerJ 12:e16505.

98. Altschul SF, Madden TL, Schaffer AA, Zhang J, Zhang Z, Miller W, Lipman DJ. 1997. Gapped BLAST and PSI-BLAST: a new generation of protein database search programs. Nucleic Acids Res 25:3389–402.

99. Altschul SF, Gish W, Miller W, Myers EW, Lipman DJ. 1990. Basic local alignment search tool. J Mol Biol 215:403–10.

100. Krogh A, Larsson B, von Heijne G, Sonnhammer EL. 2001. Predicting transmembrane protein topology with a hidden Markov model: application to complete genomes. J Mol Biol 305:567–80.

101. Paysan-Lafosse T, Andreeva A, Blum M, Chuguransky SR, Grego T, Pinto BL, Salazar GA, Bileschi ML, Llinares-Lopez F, Meng-Papaxanthos L, Colwell LJ, Grishin NV, Schaeffer RD, Clementel D, Tosatto SCE, Sonnhammer E, Wood V, Bateman A. 2025. The Pfam protein families database: embracing AI/ML. Nucleic Acids Res 53:D523–D534.

102. Lawrence M, Huber W, Pages H, Aboyoun P, Carlson M, Gentleman R, Morgan MT, Carey VJ. 2013. Software for computing and annotating genomic ranges. PLoS Comput Biol 9:e1003118.

103. Soding J, Biegert A, Lupas AN. 2005. The HHpred interactive server for protein homology detection and structure prediction. Nucleic Acids Res 33:W244–8.

104. Abramson J, Adler J, Dunger J, Evans R, Green T, Pritzel A, Ronneberger O, Willmore L, Ballard AJ, Bambrick J, Bodenstein SW, Evans DA, Hung CC, O’Neill M, Reiman D, Tunyasuvunakool K, Wu Z, Zemgulyte A, Arvaniti E, Beattie C, Bertolli O, Bridgland A, Cherepanov A, Congreve M, Cowen-Rivers AI, Cowie A, Figurnov M, Fuchs FB, Gladman H, Jain R, Khan YA, Low CMR, Perlin K, Potapenko A, Savy P, Singh S, Stecula A, Thillaisundaram A, Tong C, Yakneen S, Zhong ED, Zielinski M, Zidek A, Bapst V, Kohli P, Jaderberg M, Hassabis D, Jumper JM. 2024. Accurate structure prediction of biomolecular interactions with AlphaFold 3. Nature 630:493–500.

105. Holm L. 2020. Using Dali for Protein Structure Comparison. Methods Mol Biol 2112:29–42.

106. Kim RS, Levy Karin E, Mirdita M, Chikhi R, Steinegger M. 2025. BFVD-a large repository of predicted viral protein structures. Nucleic Acids Res 53:D340–D347.

107. van Kempen M, Kim SS, Tumescheit C, Mirdita M, Lee J, Gilchrist CLM, Soding J, Steinegger M. 2024. Fast and accurate protein structure search with Foldseek. Nat Biotechnol 42:243–246.

108. Pettersen EF, Goddard TD, Huang CC, Meng EC, Couch GS, Croll TI, Morris JH, Ferrin TE. 2021. UCSF ChimeraX: Structure visualization for researchers, educators, and developers. Protein Sci 30:70–82.

109. Gouet P, Courcelle E, Stuart DI, Metoz F. 1999. ESPript: analysis of multiple sequence alignments in PostScript. Bioinformatics 15:305–8.

