## Supplementary figures and images for "Diversity and evolution of the ribovirian class *Stelpaviricetes*"

### Supplementary material 1

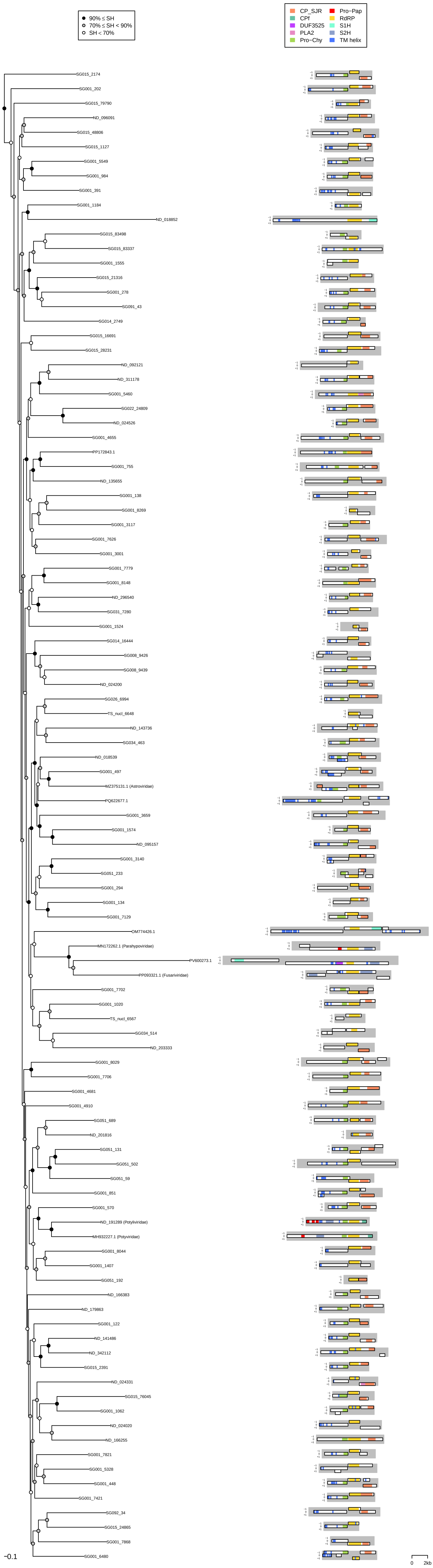

### Supplementary material 2

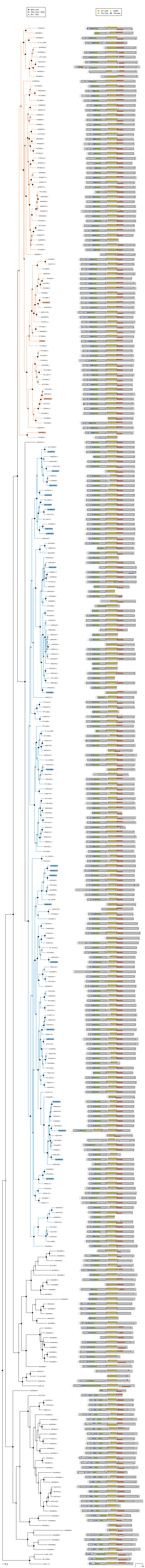

### Supplementary material 3

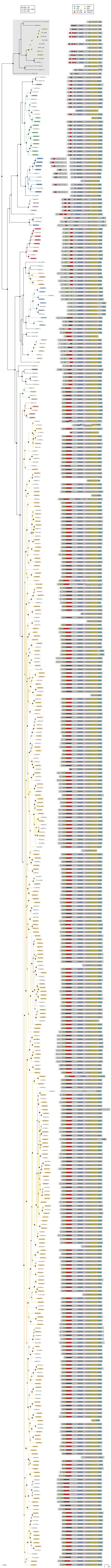

### Supplementary material 4

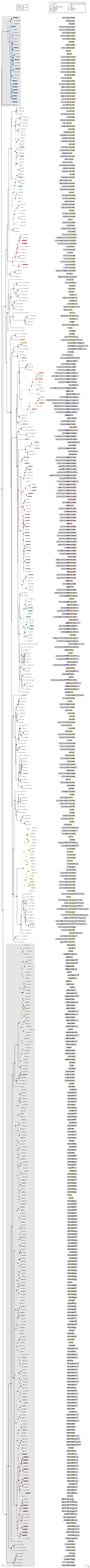
